# Condensin positioning at telomeres by shelterin proteins drives sister-telomere disjunction in anaphase

**DOI:** 10.1101/2022.03.18.484892

**Authors:** Léonard Colin, Céline Reyes, Julien Berthezene, Laetitia Maestroni, Laurent Modolo, Esther Toselli, Nicolas Chanard, Stephane Schaak, Olivier Cuvier, Yannick Gachet, Stéphane Coulon, Pascal Bernard, Sylvie Tournier

## Abstract

The localization of condensin along chromosomes is crucial for their accurate segregation in anaphase. Condensin is enriched at telomeres but how and for what purpose had remained elusive. Here we show that fission yeast condensin accumulates at telomere repeats through the balancing acts of Taz1, a core component of the shelterin complex that ensures telomeric functions, and Mit1, a nucleosome-remodeler associated with shelterin. We further show that condensin takes part in sister-telomere separation in anaphase, and that this event can be uncoupled from the prior separation of chromosome arms, implying a telomere-specific separation mechanism. Consistent with a cis-acting process, increasing or decreasing condensin occupancy specifically at telomeres modifies accordingly the efficiency of their separation in anaphase. Genetic evidence suggests that condensin promotes sister-telomere separation by counteracting cohesin. Thus, our results reveal a shelterin-based mechanism that enriches condensin at telomeres to drive in cis their separation during mitosis.

## INTRODUCTION

In eukaryotes, mitotic entry is marked by the profound reorganization of chromatin into mitotic chromosomes driven by the condensin complex (Hirano, 2016). It is acknowledged that this process, namely mitotic chromosome assembly or condensation, is an absolute pre-requisite for the accurate transmission of the genome to daughter cells, but our understanding of the mechanisms by which condensin associates with chromatin, shapes mitotic chromosomes and contributes to their accurate segregation in anaphase remains incomplete.

Condensin is a ring-shaped ATPase complex that belongs to the Structural Maintenance of Chromosomes (SMC) family of genome organizers, which also includes the cohesin complex involved in chromatin folding during interphase and in sister-chromatid cohesion (Hirano, 2016; Davidson & Peters, 2021). Condensin is composed of a core ATPase heterodimer, made of the SMC2 and SMC4 proteins, associated with a kleisin and two HEAT-repeat subunits. Most multicellular eukaryotes possess two condensin variants, named condensin I and II, made of a same SMC2/4 core but associated with distinct sets of non-SMC subunits (Ono *et al*, 2003; Hirano, 2012). Budding and fission yeasts, in contrast, possess a single condensin complex, similar to condensin I. Thereafter, condensin complexes will be collectively referred to as condensin, unless otherwise stated. There is robust evidence that condensin shapes mitotic chromosomes by massively binding to DNA upon mitotic entry and by folding chromatin into arrays of loops (Gibcus *et al*, 2018; Kakui *et al*, 2017). Thereby, condensin conceivably reduces the length of chromosomes, confers to chromosome arms the stiffness to withstand the spindle traction forces (Sun *et al*, 2018), and promotes the removal of catenations between chromosomes and sister-chromatids by orientating the activity of Topoisomerase II (Topo-II) towards decatenation (Baxter *et al*, 2011; Charbin *et al*, 2014). Hence, when condensin is impaired, sister-centromeres often reach the opposite poles of the mitotic spindle in anaphase but chromosome arms fail to separate, forming stereotypical chromatin bridges. *In vitro* studies have shown that condensin anchors itself on naked DNA through sequence-independent electrostatic interactions and uses the energy of ATP hydrolysis to extrude adjacent DNA segments into a loop of increasing size (Kschonsak *et al*, 2017; Ganji *et al*, 2018; Kong *et al*, 2020). Although such a loop extrusion reaction convincingly explains the structural properties of mitotic chromosomes (Nasmyth, 2017; Davidson & Peters, 2021), we still ignore whether and how it could take place in the context of a chromatinized genome, crowded with potential hindrances.

There is robust evidence that chromatin micro-environments impinge upon the localization of condensin. ChIP-seq studies performed on species ranging from yeasts to mammals have revealed a conserved condensin pattern along the genome, constituted of a broad and basal distribution punctuated by peaks of high occupancy at centromeres, rDNA repeats and in the vicinity of highly expressed genes (D’Ambrosio *et al*, 2008; Kim *et al*, 2013; Kranz *et al*, 2013; Dowen *et al*, 2013; Sutani *et al*, 2015). Various factors such as the chromokinesin Kif4 (Samejima *et al*, 2012), the zinc-finger protein AKAP95 (Steen *et al*, 2000), transcription-factors and chromatin remodelers (for review see (Robellet *et al*, 2017) have been involved in the binding of condensin to chromatin in yeasts or vertebrate cells. Additional cis-acting factors that increase condensin’s local concentration at centromeres and/or at rDNA repeats have been identified in budding or fission yeast (Tada *et al*, 2011; Johzuka & Horiuchi, 2009; Verzijlbergen *et al*, 2014). Such enrichments are likely to play a positive role since there is clear evidence that condensin contributes to the stiffness of centromeric chromatin and to the bilateral attachment of centromeres in early mitosis (Ono *et al*, 2004; Gerlich *et al*, 2006; Nakazawa *et al*, 2008; Ribeiro *et al*, 2009; Verzijlbergen *et al*, 2014; Piskadlo *et al*, 2017). Likewise, the segregation of the rDNA is acutely sensitive to condensin activity (Freeman *et al*, 2000; Nakazawa *et al*, 2008; Samoshkin *et al*, 2012). Highly expressed genes, in contrast, are thought to constitute a permeable barrier where active condensin complexes stall upon encounters with DNA-bound factors such as RNA polymerases (Brandão *et al*, 2019; Rivosecchi *et al*, 2021). Consistent with a local hindrance, in fission yeast, attenuating transcription that persists during mitosis improves chromosome segregation when condensin is impaired (Sutani *et al*, 2015). Further evidence in budding yeast indicates that dense arrays of protein tightly bound to DNA can constitute a barrier for DNA-translocating condensin (Guérin *et al*, 2019). Thus, depending on the context, condensin enrichment can reflect either positive or negative interplays.

Microscopy studies have clearly shown that condensin I is enriched at telomeres during mitosis and meiosis in mammalian cells (Walther *et al*, 2018; Viera *et al*, 2007), and ChIP-seq has further revealed that condensin I accumulates at telomere repeats in chicken DT40 cells, but the mechanisms underlying such enrichment as well as its functional significance have remained unknown. We and others previously showed that the separation of sister-telomeres in anaphase involves condensin regulators such as Cdc14 phosphatase in budding yeast (Clemente-Blanco *et al*, 2011), and Aurora-B kinase in fission yeast (Reyes *et al*, 2015; Berthezene *et al*, 2020), but whether and how condensin could play a role has remained unclear.

In the present study, we sought to determine how and why condensin is enriched at telomeres by using the fission yeast *Schizosaccharomyces pombe* as a model system. Telomeres contain G-rich repetitive sequences that are protected by a conserved protein complex called Shelterin (de Lange, 2018; Lim & Cech, 2021), which is composed, in fission yeast, of Taz1 (a myb-domain DNA-binding protein homologous to human TRF1 and TRF2), Rap1, Poz1 (a possible analog of TIN2), Tpz1 (an ortholog of TPP1), Pot1, and Ccq1. Whilst Taz1 binds to double-stranded G-rich telomeric repeats, Pot1 binds to 3′ single-stranded overhang. Rap1, Poz1, and Tpz1 act as a molecular bridge connecting Taz1 and Pot1 through protein–protein interactions. Ccq1 contributes to the recruitment of the nucleosome remodeler Mit1 and of telomerase (for review on fission yeast shelterin see Moser & Nakamura, 2009; Dehé & Cooper, 2010). We found that Taz1 plays the role of a cis-acting enrichment factor for condensin at telomeres, whilst Mit1 antagonized condensin’s accumulation. Thus, telomeres are a remarkable chromosomal environment where condensin is enriched by a shelterin-dependent cis-acting mechanism. Our results further indicate that the level of condensin at telomeres, set up by Taz1 and Mit1, is instrumental for their proper disjunction during anaphase, hence associating a key biological function to this local enrichment. Based on these data, we propose that condensin is enriched at telomeres *via* interplays with shelterin proteins to drive sister telomere separation in anaphase.

## RESULTS

### Fission yeast condensin is enriched at telomeric repeats during metaphase and anaphase

Fission yeast condensin, like vertebrate condensin I, is largely cytoplasmic during interphase and binds genomic DNA during mitosis (Sutani *et al*, 1999). At this stage, it shows high level of occupancy at centromeres, at rDNA repeats and in the vicinity of highly transcribed genes (Sutani *et al*, 2015; Nakazawa *et al*, 2015). However, unlike vertebrate condensin I (Kim *et al*, 2013; Walther *et al*, 2018), whether fission yeast condensin is present at telomeres had not been reported. To assess this, we performed chromatin immunoprecipitation of the kleisin subunit Cnd2 tagged with GFP (Cnd2-GFP) and analyzed the co-immunoprecipitated DNA by quantitative real time PCR (ChIP-qPCR). Figure 1A provides a reference map for the right telomere of chromosome 2 (TEL2R). *cnd2-GFP cdc2-as* shokat mutant cells were blocked at the G2/M transition and released into a synchronous mitosis. Cnd2-GFP was hardly detectable at TEL2R and along chromosome arms during the G2 arrest (Fig. 1B, t = 0 min). However, during early mitosis, Cnd2-GFP was clearly bound to telomeric repeats (the tel0 site), and to a lesser extent at more distal sites within subtelomeric elements (Fig. 1B, t = 7 min post-release). Cnd2-GFP occupancy at tel0 was in the range of the highly expressed genes *cdc22* and *exg1* used as control for enrichment (Sutani *et al*, 2015). Cnd2-GFP level further increased in anaphase (15 min post-release from the G2 block), consistent with the maximum folding of fission yeast chromosomes achieved in anaphase (Petrova *et al*, 2013) and reminiscent of the second wave of condensin binding observed during anaphase in human cells (Walther *et al*, 2018). These data show that the kleisin subunit of condensin is enriched at TEL2R during mitosis in fission yeast cells.

**Figure 1.**
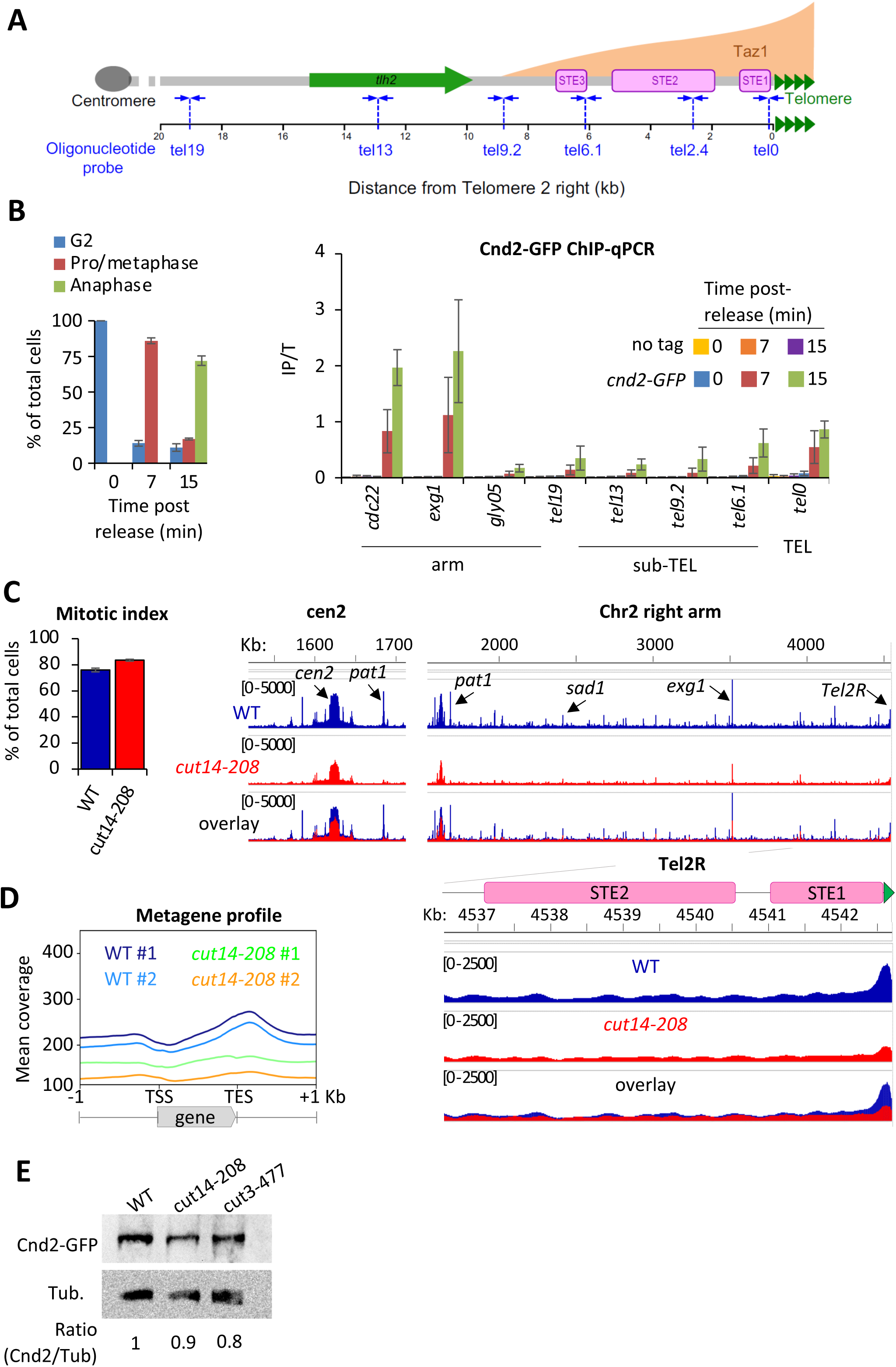
Fission yeast condensin is enriched at telomeres during metaphase and anaphase. (**A**) The telomere and sub-telomere of the right arm of chromosome 2 (Tel2R) as an example of chromosome end sequence in fission yeast. Sub-telomeric elements (STE), the heterochromatic gene *tlh2*, the domain bound by Taz1 (orange) (Kanoh *et al*, 2005) and primers for ChIP-qPCR (blue arrows) are shown. (**B**) Cnd2-GFP ChIP-qPCR from cells synchronized at G2/M (time post release 0 min) and upon their release in mitosis (time post release 7 min and 15 min). Left panel: cell cycle stages determined by scoring the accumulation of Cnd2-GFP in the nucleus (metaphase) and by DAPI staining (anaphase). Right panel: ChIP-qPCR results, *cdc22*, *exg1* and *gly05* loci, being high or low condensin binding sites which are used as controls. Shown are the averages and standard deviations (sd) from 3 independent biological and technical replicates. (**C-D**) Cnd2-GFP calibrated ChIP-seq in metaphase arrests at 36°C. (C) left panel: mitotic indexes of the two independent biological and technical replicates used. Right panel: genome browser views of replicate #1. The second is shown in Figure EV1C-D. (D) Metagene profiles of all condensin binding sites along chromosome arms from replicates #1 and #2; TSS (Transcription Start Site), TES (Transcription End Site). (**E**) Western blot showing Cnd2-GFP steady state level in indicated cells arrested in metaphase for 3h at 36°C. Tubulin (Tub.) serves as loading control.

In order to thoroughly describe condensin’s localization at telomeres, we generated calibrated ChIP-sequencing (ChIP-seq) maps of Cnd2-GFP from metaphase-arrested cells (Fig. 1 C-D). Since the current version of the fission yeast genome lacks telomere-proximal DNA and telomeric repeats, we generated a version comprising a full-length TEL2R sequence according to the described sub-telomeric and telomeric sequences (Sugawara, 1988), (Fig. S1A). Then, we measured the binding of Cnd2-GFP by calculating, at each base, the ratio of calibrated read-counts between the IP and Total (Input) fractions (Fig. S1B and Materials and Methods). As shown for centromere outer-repeats and rDNA repeats (Fig. S1C), this method allows for a better quantification of occupancy at repeated DNA sequences by correcting for biases in coverage in the Total fraction. We found Cnd2-GFP clearly enriched at telomere repeats of TEL2R in metaphase arrested cells (Fig. 1C). Cnd2-GFP binding declined rapidly over the proximal STE1 element and remained at a basal level throughout more distal elements such as STE2, STE3 and the heterochromatic *thl2* gene (Fig. 1C and Fig. S1D). To ascertain that such enrichment at telomeric repeats reflected the binding of the condensin holocomplex, we used the thermosensitive *cut14-208* and *cut3-477* mutations in the Cut14^SMC2^ and Cut3^SMC4^ ATPase subunits of condensin (Saka *et al*, 1994). Consistent with previous ChIP-qPCR data (Nakazawa *et al*, 2015), we found that the *cut14-208* mutation reduced the binding of Cnd2-GFP at centromeres (Fig. 1C), along chromosome arms (Fig. 1D), and at TEL2R (Fig. 1D and S1C). We observed similar genome-wide reduction in *cut3-477* cells, though of a smaller amplitude at TEL2R (Fig. S1E). Note that a reduction of the steady state level of Cnd2 is unlikely to explain such reductions in binding (Fig. 1E). Taken together, our data indicate that condensin accumulates at telomeric repeats during metaphase and anaphase in fission yeast.

### Condensin is required for sister-telomere disjunction in anaphase

To investigate the function of condensin at telomeres, we inactivated condensin using the thermosensitive mutations *cut14-208* or *cut3-477*, in cells whose telomeres were fluorescently labelled with Taz1-GFP. Fission yeast has three chromosomes that adopt a Rabl configuration during interphase, with telomeres clustered into 1 to 3 foci at the nuclear periphery (Chikashige *et al*, 2009; Funabiki *et al*, 1993). We previously showed that telomeres dissociate in two steps during mitosis (Reyes *et al*, 2015). In wild-type cells, the number of Taz1-GFP foci increases from 1 to up to 6 as cells transit from prophase to metaphase, i.e. when the distance between the spindle pole bodies (SPBs) increased from 0 to 4 µm (Fig. 2A, middle panel). This reflects the declustering of telomeres. During anaphase, when the distance between SPBs increases above 4 µm, the appearance of more than 6 Taz1-GFP foci indicates sister-telomere separation, and 12 foci full sister-telomere disjunction (Fig. 2A, right panel). Strikingly, *cut14-208* cells shifted to 36°C almost never showed more than six telomeric dots in anaphase, despite their centromeres being segregated at the opposite poles of the mitotic spindle (Fig. 2A and 2B). Such severe telomere dissociation defect correlates with condensin loss of function as it was not observed at the permissive temperature (25°C) (Fig. S2A). Sister-telomere disjunction was also clearly impaired in *cut3-477* mutant cells, though at milder level (Fig. 2B). To confirm the role of condensin in sister-telomere disjunction, we simultaneously visualized the behavior of LacO repeats inserted in the vicinity of telomere 1L (Tel1-GFP) together with TetO repeats inserted within centromere 3L (imr3-tdTomato) and Gar1-CFP (nucleolus) during mitotic progression (Figure S2B). After anaphase onset, as judged by the separation of sister-centromeres 3L, control cells always displayed two sister telomeric 1L foci (n=43) while 82% of *cut14-208* mutant cells (n=51) grown at non-permissive temperature remained with a single telomeric foci confirming a striking defect in the disjunction of Tel1L.

**Figure 2.**
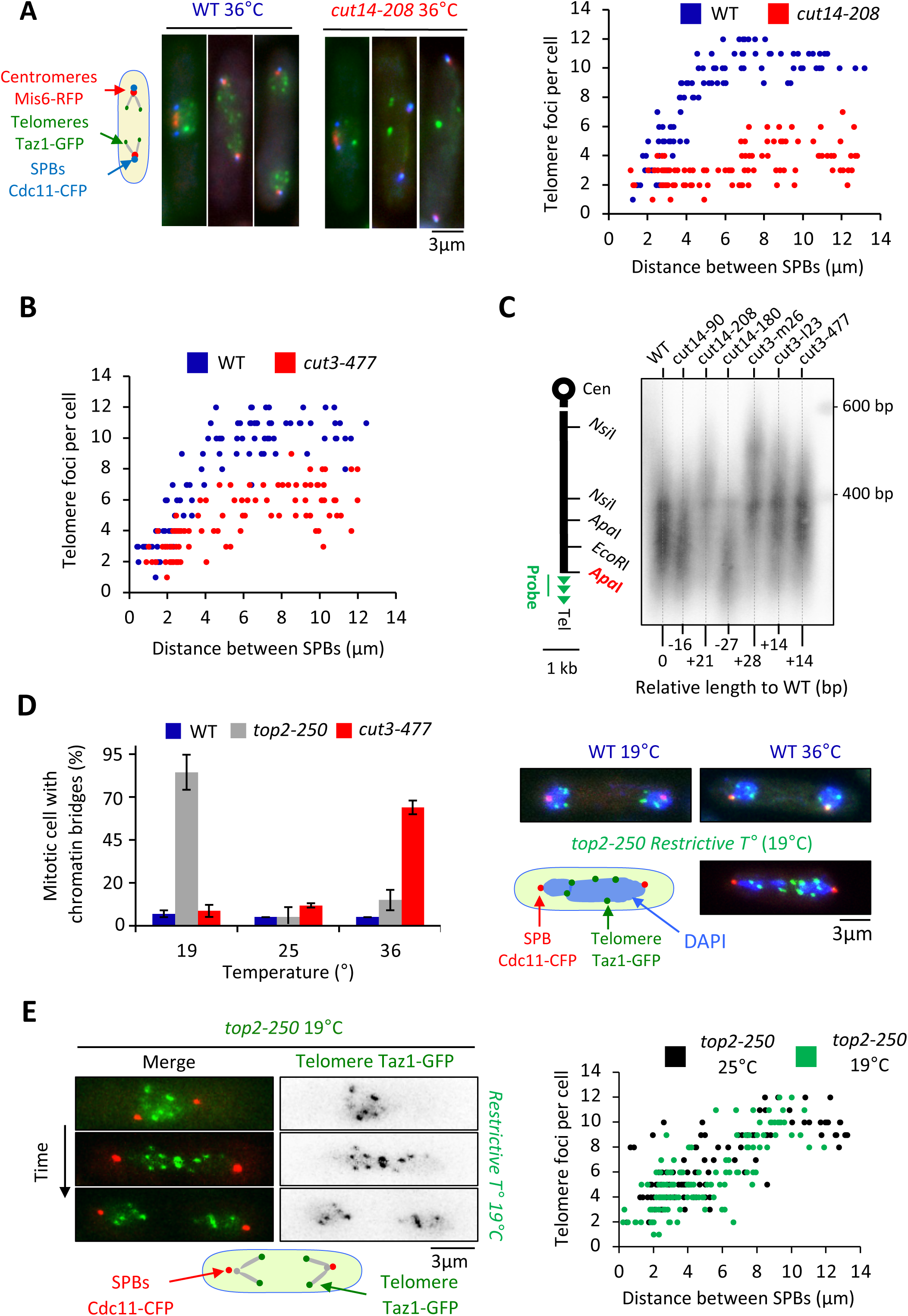
Condensin takes part in telomere disjunction during anaphase in a decatenation- independent manner. (**A**) left panel: WT or *cut14-208* condensin mutant cells shifted to the restrictive temperature of 36°C for three hours were fixed with formaldehyde and directly imaged. Telomeres were visualized via Taz1-GFP (green), kinetochores/centromeres via Mis6-RFP (red), and spindle pole bodies (SPBs) via Cdc11-CFP (blue). Right panel: number of telomeric foci according to the distance between SPBs at 36°C (n>90 cells for each strain). The data shown are from a single representative experiment out of three repeats. (**B**) Same procedure as in (A) applied to the *cut3-477* condensin mutant. (**C**) Genomic DNA from the indicated strains cultured at 32°C was digested with *Apa*I and Southern blotted using a telomeric probe (green), as represented by the grey bar. The relative gain or loss of telomeric DNA compare to WT is indicated. (**D**) Cells expressing Taz1-GFP and Cdc11-CFP, cultured at 25°C, were shifted to 19°C (restrictive temperature of *top2-250*) or 36°C (restrictive temperature of *cut3-477*), further incubated for 3 hours and fixed with formaldehyde. DNA was stained with DAPI, chromosome and telomere separation in anaphase (distance between the SPBs > 5 µm) were assessed. Shown are averages and SD obtained from three independent experiments (n=100 cells for each condition). (**E**) Left panel: Live imaging of telomere separation according to the length of the mitotic spindle (distance between the SPBs) in *top2-250* cells undergoing mitosis at 25°C or after a shift to the restrictive temperature of 19°C using fast microfluidic temperature control. Right panel: number of telomeric foci according to the distance between SPBs at 25°C or 19°C in the *top2-250* mutant. Shown is a representative experiment out of three replicates with n>70 cells, each.

Next, we wondered whether a change in telomere length could cause telomere disjunction defects as described previously (Miller & Cooper, 2003). We measured telomere length in various condensin mutant cells including *cut14-208* and *cut3-477* mutants. We only observed little variation of telomere length (Fig. 2C), unlikely to be responsible for the failure to disjoin sister- telomeres when condensin is impaired. An alternative and more likely possibility was that persistent entanglements left between chromosome arms upon condensin loss of function prevented the transmission of traction forces from centromeres to telomeres in anaphase. To test this hypothesis, we assessed telomere disjunction in cold-sensitive *top2-250* mutant cells, whose Topo II decatenation activity becomes undetectable at 20°C (Uemura *et al*, 1987). As expected, *top2-250* cells cultured at the restrictive temperature exhibited frequent chromatin bridges during anaphase (Fig. 2D, compare *top2-250* at 19°C with *cut3-477* at 36°C), and lagging centromeres (Fig. S2B). Yet, and remarkably, telomere disjunction remained effective during anaphase at 19°C, even within chromatin bridges (Fig. 2D-E). The decatenation activity of Topo II and the full separation of chromosome arms are therefore largely dispensable for telomere disjunction. Thus, these results indicate (1) that the function of condensin in sister-telomere separation is mostly independent of Topo II decatenation activity, and (2) that the separation of chromosome arms is not a prerequisite for the disjunction of sister-telomeres. Condensin might therefore play a specific role at telomeres for their proper separation during anaphase.

### Condensin takes part in the declustering of telomeres during early mitosis

To assess whether condensin might shape telomere organization prior to anaphase, we generated Hi-C maps of cells arrested in metaphase (Fig. 3A). As previously reported (Kakui *et al*, 2017), we observed frequent centromere-to-centromere and telomere-to-telomere contacts between the three chromosomes in wild-type cells (Fig. 3B). In the *cut14-208* condensin mutant at restrictive temperature, contact frequencies within chromosome arms were reduced in the range of 100 kb to 1 Mb (Fig. 3C-D), as expected from an impaired mitotic-chromosome folding activity. In contrast, contacts frequencies between telomeres were increased, both within chromosomes (intra) and in-between chromosomes (inter) (Fig. 3D). Contacts frequencies between centromeres exhibited no significant change. Aggregating Hi-C signals at chromosome ends further revealed that intra-chromosomal contacts dominate over inter-chromosomal contacts in wild-type cells and were increased in the mutant (Fig. 3E, see material and methods). Similar results obtained from a second biological and technical replicate are shown in Figure S3. Taken together these data suggest that fission yeast chromosomes enter mitosis in a Rabl configuration, with telomeres clustered together and that condensin drives their declustering into pairs of sister-telomeres as cells progress towards metaphase (Fig. 3 and S3) and their full-separation in anaphase (Fig. 2).

**Figure 3.**
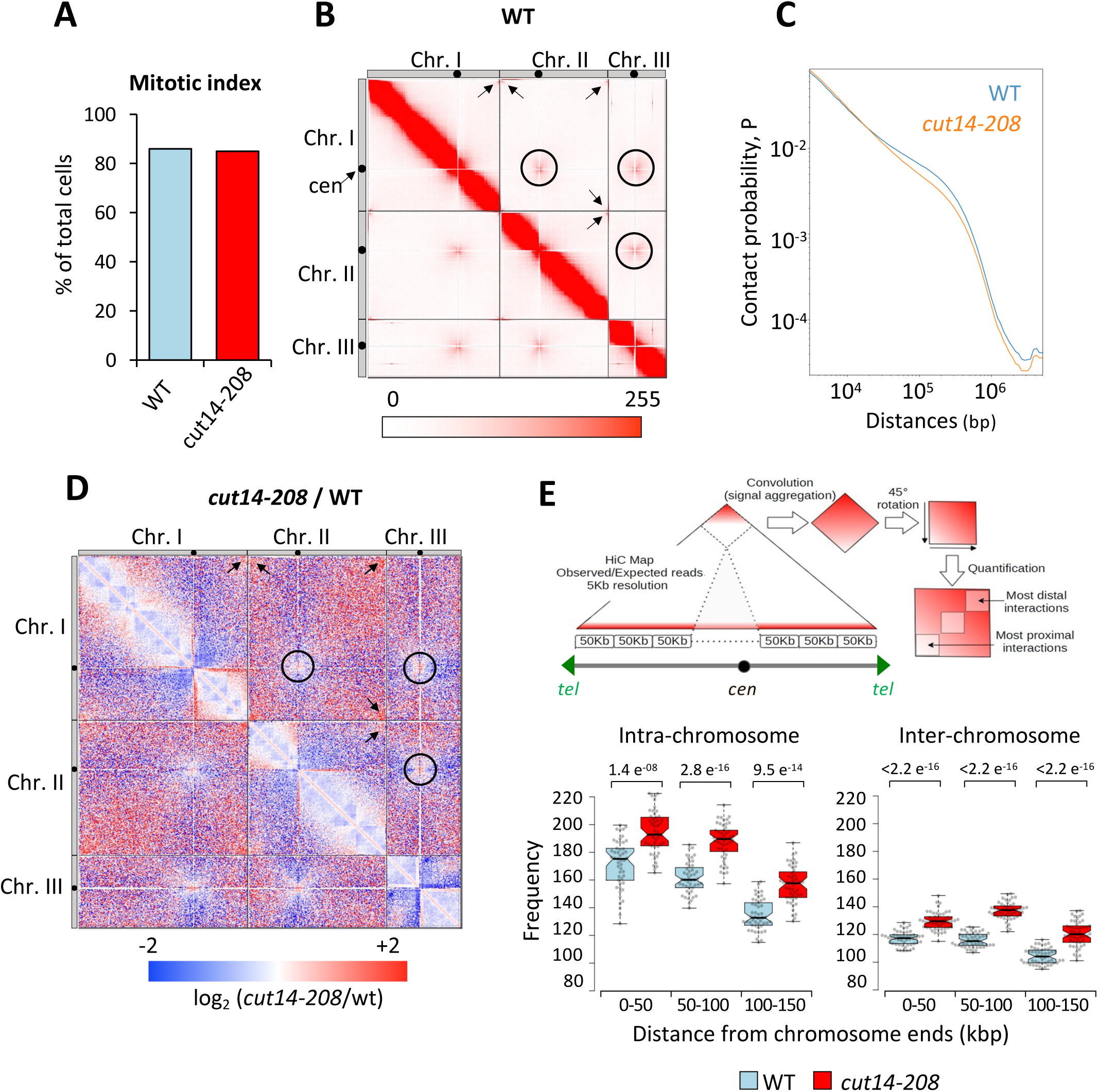
Condensin deficiency increases contact frequencies between telomeres in metaphase. (**A**) Mitotic indexes of cell cultures used for Hi-C. (**B**) Hi-C contact probability matrix at 25 kb resolution of wild-type metaphase arrests at 33°C. Contacts between telomeres (arrows) and centromeres (circles) are indicated. (**C**) Median contact probabilities as a function of distance along chromosomes for wild-type and *cut14-208* metaphases at 33°C. (**D**) Differential Hi-C contact map between wild-type and *cut14-208*. (**E**) Measurements of aggregated contact frequencies at high resolution (5 kb) over the ends of chromosomes in metaphase arrests at 33°C. Boxes indicate the median, 1st and 3rd quartiles, whiskers the minimum and maximum, and notches represent the 95% confidence interval for each median. Data points are shown as grey circles. The significance in contact frequencies was confirmed statistically by Mann-Whitney- Wilcoxon test between *cut14-208* and wild-type conditions.

### Condensin enrichment at telomeres result from positive and negative interplays with telomeric proteins

To further investigate how condensin takes part in telomere disjunction in anaphase, we sought for a cis-acting factor controlling condensin localisation specifically at telomeres. We first considered the shelterin complex and assessed Cnd2-GFP binding by calibrated ChIP-qPCR in *taz1*Δ or *rap1*Δ cells arrested in metaphase (Fig. 4A and Material and Methods). In cells lacking Taz1, Cnd2-GFP occupancy was reduced almost twofold at telomeres (tel0 site) and sub-telomeres (tel2.4 site), while remaining unchanged at the kinetochore and within chromosome arms. In contrast, the *rap1*Δ mutant showed no change compared to wild-type. A different normalization method produced similar results (Fig. S4A). Since telomere size is increased to similar extents in *taz1*Δ and *rap1*Δ mutants (Cooper *et al*, 1997; Miller *et al*, 2005), it is unlikely that condensin is titrated out from the tel0 site by supernumerary telomeric repeats in *taz1*Δ cells. Taz1 directly binds to telomeric repeats but also to non-repeated DNA motifs within chromosome arms (Zofall *et al*, 2016; Toteva *et al*, 2017). However, the binding of Cnd2-GFP was basal and independent of Taz1 at such non-telomeric Taz1-islands (Fig.4B-C and Fig. S4B-C), suggesting that Taz1 is unlikely to directly recruit condensin onto chromosomes. In line with this, we observed no physical interaction between condensin and Taz1, either by co-IP or by yeast two-hybrid assay (our unpublished data). Thus, the density of Taz1 binding sites and/or the telomeric context might be instrumental for locally enriching condensin. We therefore conclude that the core shelterin protein Taz1 plays the role of a cis-acting enrichment factor for condensin at telomeres.

**Figure 4.**
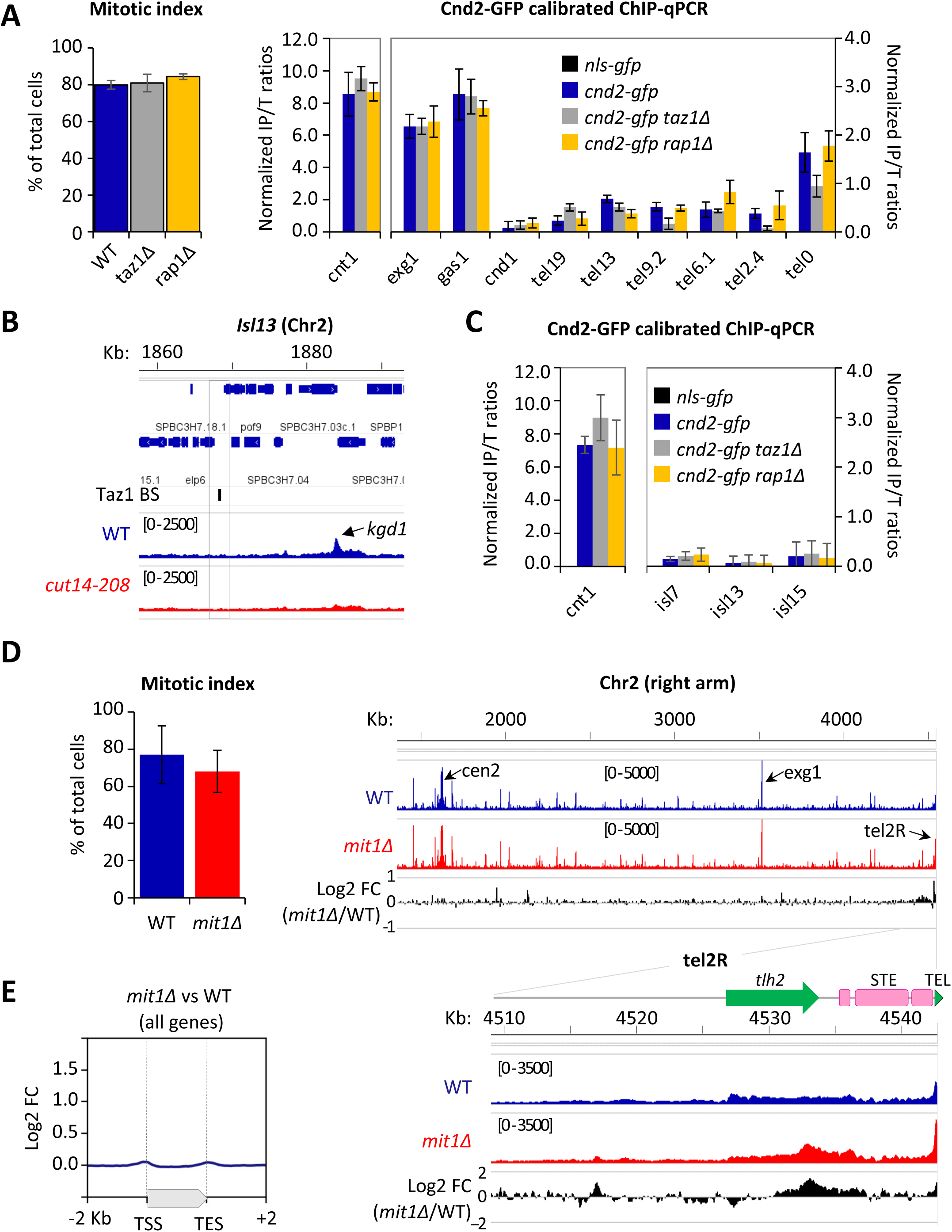
Condensin enrichment at telomeres results from positive and negative interplays with telomeric factors. (**A**) Cnd2-GFP calibrated ChIP-qPCR from cells arrested in metaphase at 30°C. Shown are averages and standard deviations (SD) of mitotic indexes and ChIP-qPCRs for 3 biological and technical replicates. *cnt1* is the kinetochore domain of *cen1*, *exg1*, *gas1* and *cnd1* are high or low occupancy binding sites on chromosome arms. (**B-C**) Cnd2-GFP occupancy assessed at non-telomeric Taz1 islands (isl) in the same samples as in Figure 1C & S1-D and Figure 4A, respectively. (**D-E**) Cnd2-GFP calibrated ChIP-seq in metaphase arrests. (D) left panel: mitotic indexes of the two independent biological and technical replicates used. Right panel: genome browser views of replicate #1. The second replicate is shown in Figure S4. (E) Metagene profiles of all condensin binding sites along chromosome arms from replicates #1 and #2; TSS (Transcription Start Site), TES (Transcription End Site).

The ATP-dependent chromatin remodeler Mit1 was another telomeric factor of interest. Indeed, Mit1 maintains nucleosome occupancy through its association with the shelterin and *mit1*Δ cells show a reduced histone H3 occupancy at sub-telomeres (van Emden *et al*, 2019). Since we previously reported that nucleosome eviction underlies condensin’s binding to chromosomes (Toselli-Mollereau *et al*, 2016), we assessed condensin binding at telomeres in cells lacking Mit1. As expected, we observed an increased condensin occupancy at telomeres and sub-telomeres by calibrated-ChIP-seq (Fig. 4D) and ChIP-qPCR (Fig. S4-D). ChIP-seq further showed that such increase was largely, if not strictly, restricted to chromosome ends (Fig. 4D-E and Fig. S4E). These data strongly suggest that Mit1 counteracts condensin localization at telomeres. Thus, taken together, our results suggest that the steady state level of association of condensin with telomeres results from the balancing acts of shelterin proteins and associated factors, amongst which Taz1 and Mit1.

### Condensin acts in *cis* to promote telomere disjunction in anaphase

Since the *cut14-208* and *cut3-477* mutations reduce condensin binding all along chromosomes, it was difficult to ascertain the origin of the telomere disjunction defect in these mutants. However, the finding that condensin localization at telomeres partly relies on Taz1 and Mit1 provided a means to assess whether condensin could drive telomere disjunction in *cis*. If it were the case, then removing Taz1 in a sensitized *cut3-477* background, to further dampen condensin at telomeres, should strongly increase the frequency of sister-telomere non-disjunctions compared to single mutants. Conversely, removing Mit1 in *cut3-477* cells should rescue sister-telomere disjunctions. We observed very few non-disjunction events during anaphases in *taz1*Δ single mutant cells at 32°C (Fig. 5A). We speculate that the residual amount of condensin that persists at telomeres when Taz1 is lacking might be sufficient to ensure their efficient disjunction. However, combining *cut3- 477* and *taz1*Δ caused a synergistic increase of the frequency of sister-telomere non-disjunction (Fig. 5A) that correlated with a synthetic negative growth defect at 32°C and 34°C (Fig. 5B). Conversely, eliminating Mit1 rescued sister-telomere disjunction in *cut3-477* mutant cells (Fig. 5C). Taken together, these data indicate that the level of condensin bound to telomeres is a limiting parameter for their efficient separation in anaphase, suggesting therefore that condensin controls sister-telomeres disjunction in *cis*.

**Figure 5.**
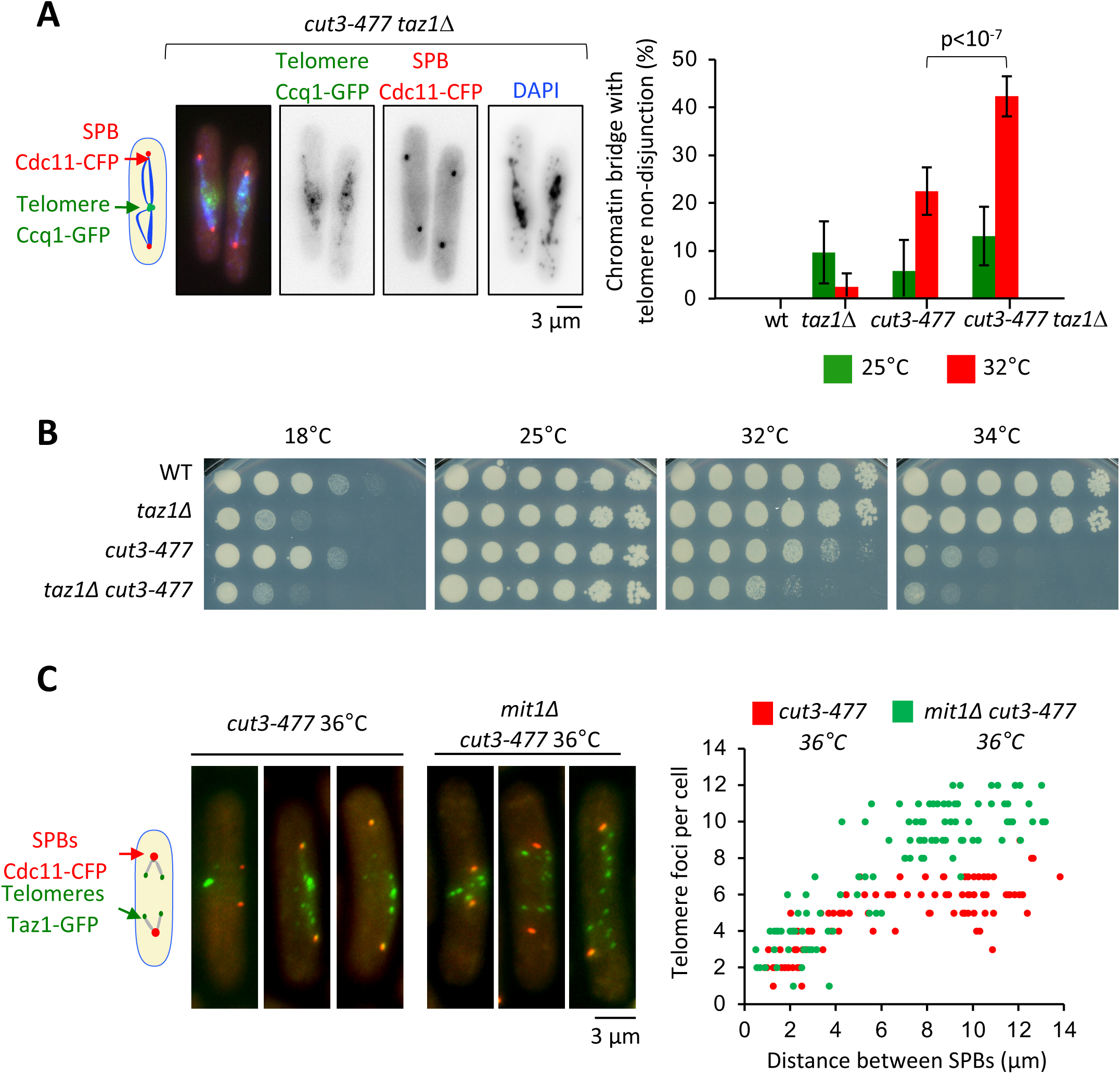
Condensin level at telomeres is a limiting parameter for their disjunction during anaphase. (**A**) Cells were grown at 25°C or shifted to 32°C for 3 hours, fixed with formaldehyde and stained with DAPI to reveal DNA. Left panel: example of anaphase cells showing chromatin bridges and non-disjoined telomeres in late anaphase in the *cut3-477 taz1*Δ double mutant. Right panel: telomere non-disjunction events were scored in anaphase cells. Shown are averages and standard deviation from 3 independent biological and technical replicates with n=100 cells, each. (**B**) Cells of indicated genotypes were serially diluted 1/5 and spotted onto YES plates at indicated temperatures for 7 (18°C), 3 (25°C) and 2 (32 and 34°C) days. (**C**) Left panel: *cut3-477* or *cut3- 477 mit1*Δ mutant cells shifted to the restrictive temperature of 36°C for three hours were fixed with formaldehyde and directly imaged. Telomeres were visualized via Taz1-GFP (green) and spindle pole bodies (SPBs) via Cdc11-CFP (blue). Right panel: number of telomeric foci according to the distance between SPBs at 36°C (n>90 cells for each strain). The data shown are from a single representative experiment out of three repeats.

### Condensin counteracts cohesin at telomeres

We previously showed that eliminating the heterochromatin protein Swi6^HP1^ alleviates the telomere separation defect caused by the inhibition of Ark1 (Reyes *et al*, 2015). Since Ark1 controls condensin association with chromosomes (Petersen & Hagan, 2003; Tada *et al*, 2011), and Swi6 enriches cohesin at heterochromatin domains, including telomeres (Bernard *et al*, 2001), we asked whether interplays between condensin and cohesin might underlie telomere separation during anaphase. To test this, we assessed the impact of the cohesin mutation *rad21-K1*, known to weaken sister-chromatid cohesion (Bernard *et al*, 2001), on telomere disjunction. First, we observed that sister-telomere separation occurs at a smaller mitotic spindle size in the *rad21-K1* mutant as compared to wild-type, indicating an accelerated kinetics during mitosis (Fig. 6A). Second, weakening cohesin partly rescued telomere disjunction when condensin was impaired, as suggested by the increased number of telomeric dots displayed by *cut3-477 rad21-K1* double mutant cells in anaphase (Fig. 6A). A similar rescue was observed when Rad21 was inactivated in early G2 cells purified using a lactose gradient (Fig. S6), indicating that cohesin inactivation post cohesion establishment complemented the telomere disjunction defects of a condensin mutant. These observations indicate that cohesin hinders the separation of sister-telomeres, suggesting therefore that condensin might counteract cohesin at telomeres. To test this hypothesis, we assessed cohesin binding to chromosomes in the *cut3-477* condensin mutant. Cells were arrested at the G2/M transition, shifted to the restrictive temperature to inactivate condensin while maintaining the arrest, and released into a synchronous mitosis (Fig. 6B). Cohesin binding was assessed by calibrated ChIP-qPCR against the Psm3^SMC3^ subunit of cohesin tagged with GFP (Psm3-GFP). We observed no strong change in cohesin occupancy between G2 and anaphase in wild-type cells, consistent with the idea that solely 5-10% of the cohesin pool is cleaved by separase at the metaphase to anaphase transition in fission yeast (Tomonaga *et al*, 2000). In *cut3- 477* mutant cells, however, we observed a strong increase in Psm3-GFP levels at telomeres (tel0 site) and sub-telomeres (tel1.2 site), but no change at further distal sites, nor within chromosome arms or at centromeres (Fig. 6B and S5). This specific increase in occupancy at telomeres and sub- telomeres in the condensin mutant was readily visible both during G2 and anaphase. Altogether, our data indicate that cohesin restrain telomere disjunction in anaphase and that condensin prevents the accumulation of cohesin at telomeres.

**Figure 6.**
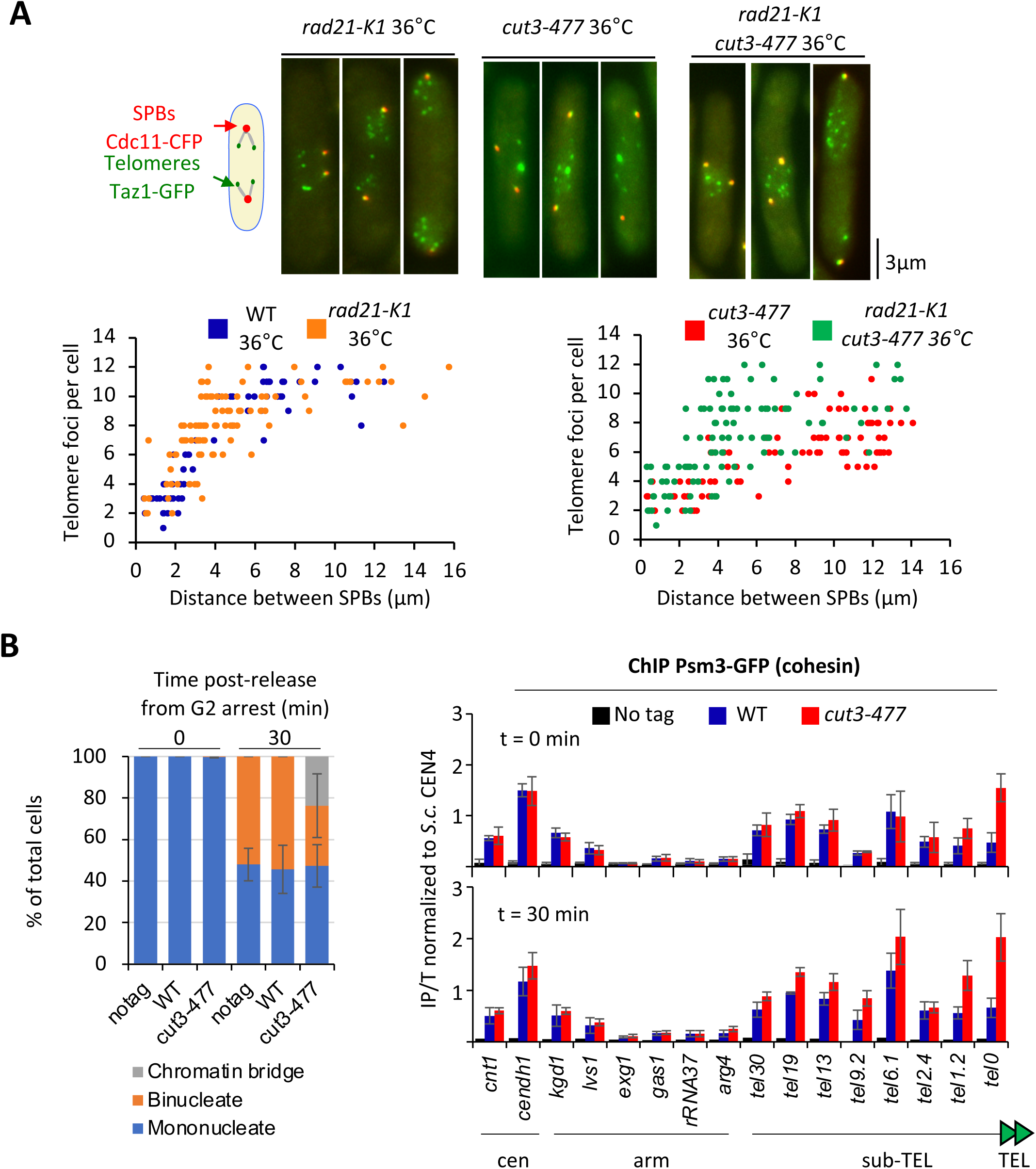
Condensin counteracts cohesin at telomeres. **(A)** Top panel: WT, *rad21-K1* or *rad21- K1 cut3-477* cells were shifted to the restrictive temperature of 36°C for three hours, fixed with formaldehyde and directly imaged. Telomeres were visualized via Taz1-GFP (green) and spindle pole bodies (SPBs) via Cdc11-CFP (blue). Lower panels: number of telomeric foci according to the distance between SPBs at 36°C (n>90 cells for each strain). The data shown are from a single representative experiment out of three repeats. (**B**) Psm3-GFP calibrated ChIP-qPCR from cells synchronized in G2/M and shifted at 36°C to inactivate condensin during the G2 block (time 0 min) and upon their release in anaphase (time 30 min). Left panel: cell cycle stages determined by DAPI staining. Right panel: ChIP-qPCR results. *cendh1*, *kgd1* and *lvs1* are cohesin binding sites, while *exg1*, *gas1* and *rRNA37* are condensin binding sites. Percentage of IP with Psm3-GFP have been normalized using *S. cerevisiae* (S.c.) CEN4 locus. Shown are the averages and standard deviations from 3 independent biological and technical replicates.

## DISCUSSION

With this work, we show that condensin is enriched at fission yeast telomeres during mitosis and that such enrichment results from the balancing acts of telomeric proteins. We also show that the separation of sister-telomeres is not the mere consequence of the separation of chromosome arms and that condensin acts in cis at telomeres to drive their disjunction during anaphase. We provide evidence that condensin might achieve this task by counteracting cohesin.

Previous work has shown that the kleisin subunit of condensin II (CAPH2) binds human TRF1, a counterpart of Taz1, in RPE-1 cells (Wallace *et al*, 2019), but since no physical or functional link has been described between TRF1 and other subunits of the condensin II holocomplex, it was unclear whether CAPH2 might act at telomeres independently of condensin II. Hence, the biological significance of the presence of condensin complexes at telomeres remained enigmatic.

Here we show that the kleisin subunit of fission yeast condensin is bound to the telomere repeats of TEL2R in metaphase and anaphase and that such binding relies on the Cut14^SMC2^ and Cut3^SMC4^ ATPases (Saka *et al*, 1994), arguing therefore that the condensin holocomplex is bound to TEL2R. Since southern blotting and FISH experiments have shown that chromosome I and II contains similar sub-telomeric elements (Funabiki *et al*, 1993; Oizumi *et al*, 2021), our observations made using TEL2R DNA are likely to be relevant to most fission yeast telomeres.

We further show that condensin occupancy at telomeres is controlled by the telomeric proteins Taz1 and Mit1. Taz1 being a core component of the shelterin complex, we cannot formally rule out that the reduced binding of condensin stems from a collapse of the overall telomeric structure in cells lacking Taz1. However, two observations argue against such scenario. First, the fact that deleting Rap1, another key component of shelterin, does not impair condensin localisation and, second, our finding that condensin binding to telomeres is also controlled, negatively, by the nucleosome remodeler Mit1, which associates with telomeres *via* the Ccq1 subunit of shelterin (Sugiyama *et al*, 2007; van Emden *et al*, 2019). Such negative regulation by Mit1 not only strengthens our previous work suggesting that nucleosome arrays are an obstacle for condensin binding to DNA *in vivo* (Toselli-Mollereau *et al*, 2016), but also strongly suggests that condensin localisation at telomeres relies on a dedicated pathway that involves interplays with telomeric components. However, and in sharp contrast with the human TRF1 and CAP-H2 (Wallace *et al*, 2019), we detected no protein-to-protein interactions between Taz1 and Cnd2/condensin. Together with our observation that Taz1 does not enrich condensin at discrete Taz1-DNA binding sites located outside telomeres, this suggests that Taz1 is not a cis-acting recruiter for condensin at telomeres. Rather, by analogy with the accumulation of condensin at highly expressed genes (Brandão *et al*, 2019; Rivosecchi *et al*, 2021), we speculate that arrays of Taz1 proteins tightly bound to telomere repeats might create a permeable barrier onto which condensin molecules accumulate. However, unlike highly expressed genes that most likely hinder condensin-mediated chromosome segregation in anaphase (Sutani *et al*, 2015), the Taz1 barrier would play a positive role in chromosome segregation by promoting sister-telomere disjunction in anaphase.

Using Hi-C and live cell imaging, we provide evidence that condensin takes part in telomere declustering during the early steps of mitosis and in sister-telomere disjunction in anaphase. It is tempting to speculate that condensin promotes the dissociation of telomeric clusters, inherited from the Rabl organisation of chromosomes in interphase, by folding chromatin into mitotic chromosomes. As condensation proceeds, axial shortening and stiffening of chromosome arms would drive the movement of the pairs of telomeres located at the opposite ends of a chromosome away from each other. The separation of sister-telomeres during anaphase, in contrast, cannot be the passive consequence of the separation of sister chromatids. Indeed, the striking observation that sister-telomere disjunction can be uncoupled from the separation of chromosome arms, as seen in the decatenation-defective *topo-250* mutant, implies the existence of a mechanism independent of chromosome arms, driving sister-telomere disjunction. In fission yeast, condensin must play a key role within such telomere-disjunction pathway (TDP) since modulating its occupancy at telomeres whilst leaving chromosome arms largely unchanged, using *taz1*Δ or *mit1*Δ mutations, is sufficient to change accordingly the efficiency of sister-telomeres disjunction. The fact that condensin occupancy at telomeres is a limiting parameter for their disjunction argues for a role played in *cis*. This finding is reminiscent of telomere separation in human cells that specifically relies on the activity of the poly(ADP-ribose) polymerase tankyrase 1 (Dynek & Smith, 2004), and suggest therefore that the existence of a dedicated pathway for sister-telomere disjunction is a conserved feature of eukaryotic cells. We therefore conclude that condensin enriched at telomeres *via* the balancing acts of Taz1 and Mit1 drives the separation of sister-telomeres in anaphase. The corollary is that failures to disjoin sister-telomeres most likely contribute to the stereotypical chromatin bridge phenotype exhibited by condensin-defective cells. Our results do not rule out the possibility that Topo II contributes to telomeres disentanglements, but nevertheless imply that Topo II catalytic activity is dispensable for telomere segregation provided that condensin is active. The close proximity of DNA ends could explain such a dispensability. It has been reported in budding yeast that the segregation of LacO repeats inserted in the vicinity of TelV is impaired by the *top2-4* mutation (Bhalla *et al*, 2002). At first sight, this appears at odds with our observations made using the telomere protein Taz1 tagged with GFP. However, since LacO arrays tightly bound by LacI proteins constitute a barrier for the recoiling activity of budding yeast condensin in anaphase (Guérin *et al*, 2019), the insertion of such a construct might have created an experimental condition in which condensin activity was specifically impaired at TELV, hence revealing the contribution of Topo II. In addition, the telomere structure in budding and fission yeast is significantly different. Budding yeast protects its telomeres *via* two independent factors, Rap1 and the Cdc13-Stn1-Ten1 complex, whereas in fission yeast Taz1 and Pot1 are bridged by a complex protein interaction network (Rap1-Poz1-Tpz1). This is a remarkable conserved structural feature between the shelterin of *S. pombe* and the human shelterin. Notably, it was recently shown that the telomeric components of *S. pombe* can dimerize leading to a higher complex organization of the shelterin (Sun *et al*, 2022). It is thus likely that dimerization of Taz1, Poz1, and the Tpz1-Ccq1 subcomplex may also contribute to the clustering of sister and non-sister chromatid telomeres. The architectural differences in telomere organization between budding and fission yeast may require different mechanisms to properly segregate telomeres during mitosis.

Understanding how condensin takes part in the disjunction of sister-telomeres will require identifying the ties that link them. Cohesin has been involved in telomere cohesion in budding yeast (Antoniacci & Skibbens, 2006; Renshaw *et al*, 2010), but in human cells the situation remains unclear. Although Scc3^SA1^ is a likely target of the tankyrase 1 pathway for telomere disjunction (Canudas & Smith, 2009), telomeric cohesion appears independent of other cohesin subunits (Bisht *et al*, 2013). Our finding that *rad21-K1*, a loss-of-function mutation in the kleisin subunit of fission yeast cohesin, accelerates sister-telomere disjunction in an otherwise wild-type genetic background would be consistent with a role for cohesin in ensuring cohesion between sister- telomeres in fission yeast. Alternatively, *rad21-K1* might indirectly increase the impact of condensin at chromosome ends, for instance by altering the structure of sub-telomeric heterochromatin (Dheur *et al*, 2011). However, such an indirect effect seems less likely because the kinetics of sister-telomere disjunction is not accelerated in cells lacking the core heterochromatin protein Swi6 (Reyes *et al*, 2015). Therefore, we favour the conclusion that condensin drive sister-telomere disjunction by counteracting cohesin at chromosome ends. Whether it could be cohesive- or loop-extruding- cohesin remains to be determined, but we note that an antagonism between condensin and cohesin for the folding of interphase chromatin as well as for telomere segregation in anaphase has been reported in Drosophila and budding yeast, respectively (Rowley *et al*, 2019; Renshaw et al. 2010). Thus, unravelling the mechanism by which condensin drives telomere disjunction in anaphase will require further investigations not only on the interplays between condensin and cohesin at telomeres, but also on the role played or not by condensin loop-extrusion activity and on the dynamics of shelterin. Because of its ability to organize telomeres into various structures (Lim & Cech, 2021), the shelterin complex may link sister-telomeres together and loop extrusion by condensin may provide the power stroke to disentangle such structures. Thus, as speculated in Figure 7, the accumulation of condensin against Taz1^TRF1^ barriers, together with a possible up-regulation by Aurora-B kinase in anaphase (Reyes *et al*, 2015), might allow condensin-mediated DNA translocation to pass a threshold beyond which the ties between sister-telomeres would be eliminated, be it cohesin- and/or shelterin-mediated. Whatever the mechanism, given the conservation of shelterin, and the abundance of condensin complexes at telomeres during mitosis and meiosis in mammals (Viera *et al*, 2007; Walther *et al*, 2018), we speculate that condensin specifically drives the separation of telomeres in other living organisms.

**Figure 7.**
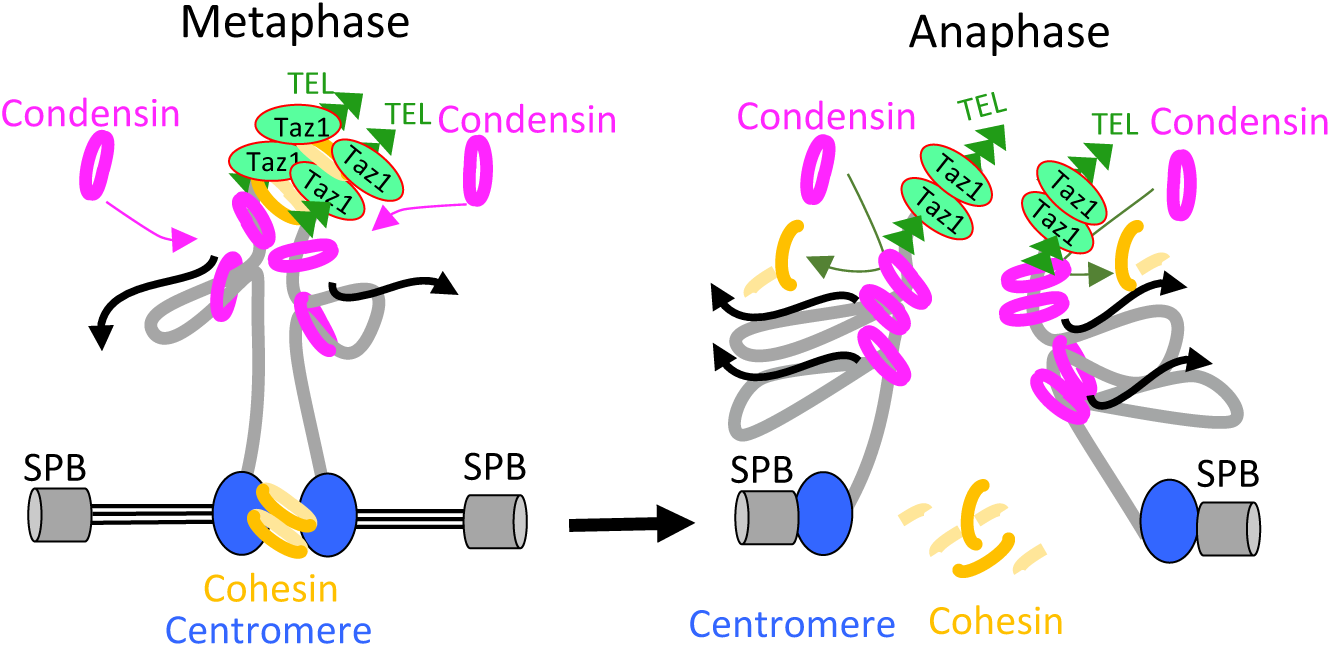
Model for condensin-driven sister-telomere disjunction. Loop extruding condensin accumulates against a barrier formed by arrays of Taz1 proteins bound to telomeric repeats, allowing condensin-mediated DNA translocation to pass a threshold beyond which the ties between sister-telomeres such as cohesin would be eliminated. See Discussion for details.

## ACKNOWLEDGMENTS

We thank the National BioResource-Yeast Project, J. Cooper, JP. Javerzat, M. Yanagida and J. Kanoh for strains; J. Baxter and N. Minchell for teaching the rudiments of Hi-C to L.C., J.P. Javerzat, F. Beckouet, V. Vanoosthuyse and A. Piazza for helpful discussions.

## AUTHOR CONTRIBUTIONS

Conceptualization and Methodology: S.T.; P.B., S.C., Y.G.

Investigation: all authors

Visualization: L.C., C.R., J.B, S.T., P.B., S.C., Y.G.

Funding acquisition: S.T., P.B., S.C., Y.G.

Project administration: S.T., P.B.

Supervision: S.T., P.B., S.C., Y.G., O.C.

Writing – original draft: S.T., P.B.

Writing – review & editing: S.T., P.B., S.C., Y.G., O.C.

## SUPPLEMENTARY DATA

**Figure S1.**
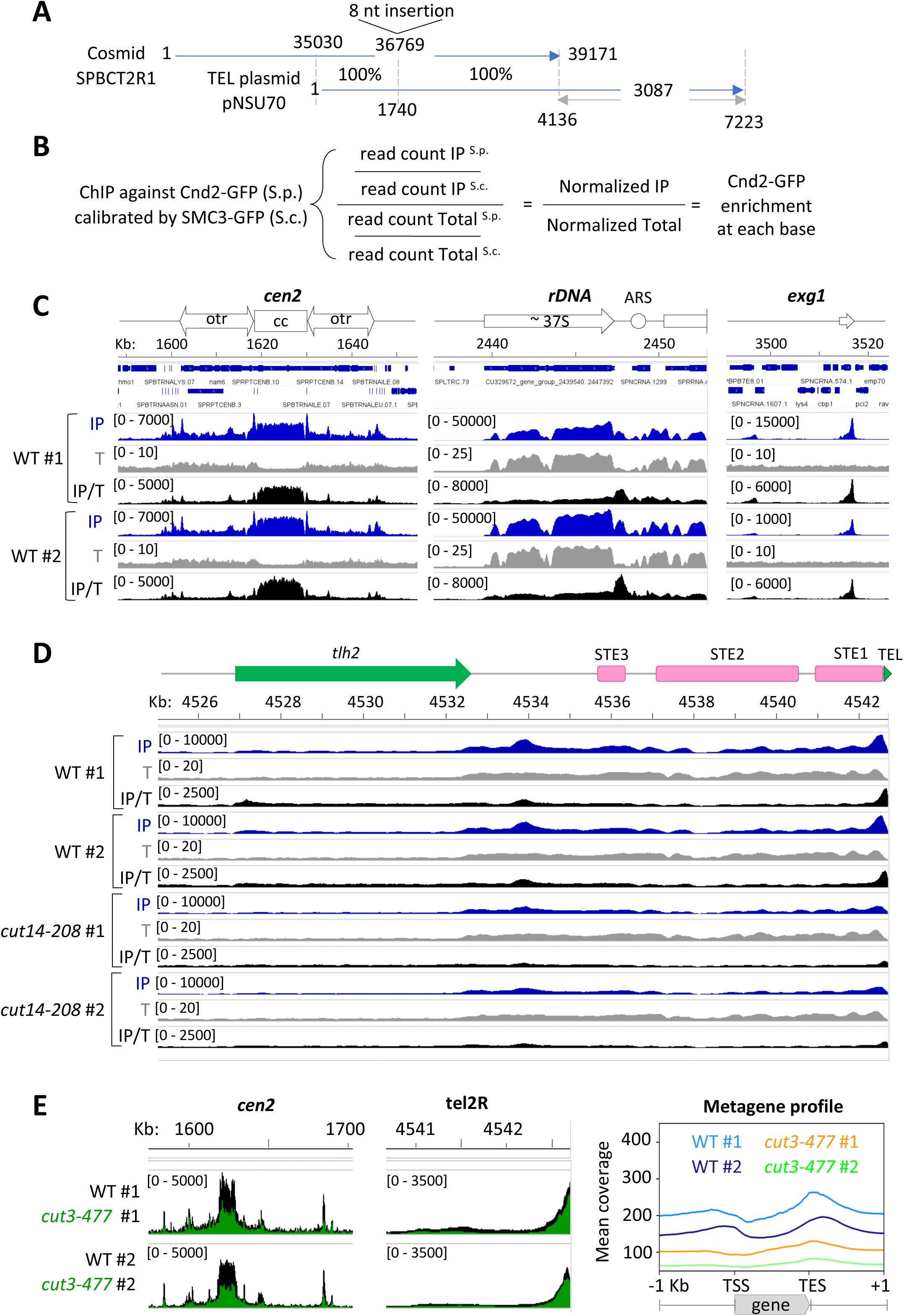
Fission yeast condensin is enriched at telomeres during metaphase and anaphase. **(A)** Telomeric plasmid pNSU70 (Sugawara, 1988) aligned against the *S. pombe* genome (ASM294v2) using Blast shows a best hit with cosmid SPBCT2R1 that corresponds to the right end of chromosome 2 (nucleotides 4500619 to 4539800 in the ASM294v2 genome). SPBCT2R1 was aligned with pNSU70 using Clustal Omega (Madeira *et al*, 2019) and the nucleotides 4137 to 7223 of pNSU70 were added to the sequence of chromosome 2 at position 4539800 to create a TEL2R-extended version of the genome, which is available under the accession number GSE196149. **(B)** Principle of the base per base normalisation method applied to ChIPs to assess the occupancy of Cnd2-GFP from *S. pombe* (S.p.) at repeated DNA elements using chromatin from *S. cerevisiae* (S.c.) cells expressing SMC3-GFP for internal calibration. (**C**) Results of the base per base normalization method applied to calibrated ChIP-seq data to correct for biases in coverage in the IP and Total (T) fractions. Calibrated read counts obtained in the IP and Total (T) fractions, and their base per base ratios (IP/T) obtained at centromere 2 (*cen 2*), composed of a central core (cc) flanked by repetitive heterochromatic outer repeats (otr) and at rDNA repeats are shown. The single-copy gene *exg1* serves as a non-repetitive control. (**D**) Calibrated read-counts and base per base IP/T ratios obtained at TEL2R for two biological and technical replicates. Data from replicate #1 are also shown in Figure 1C. (**E**) Results from calibrated ChIP-seq against Cnd2-GFP in metaphase arrests WT and *cut3-477* condensin mutant. Shown are the base per base ratios and metagene profiles obtained from two biological and technical replicates. Mitotic indexes of cell cultures were: WT (77% ± 6%), *cut3-477* (70% ± 5%).

**Figure S2.**
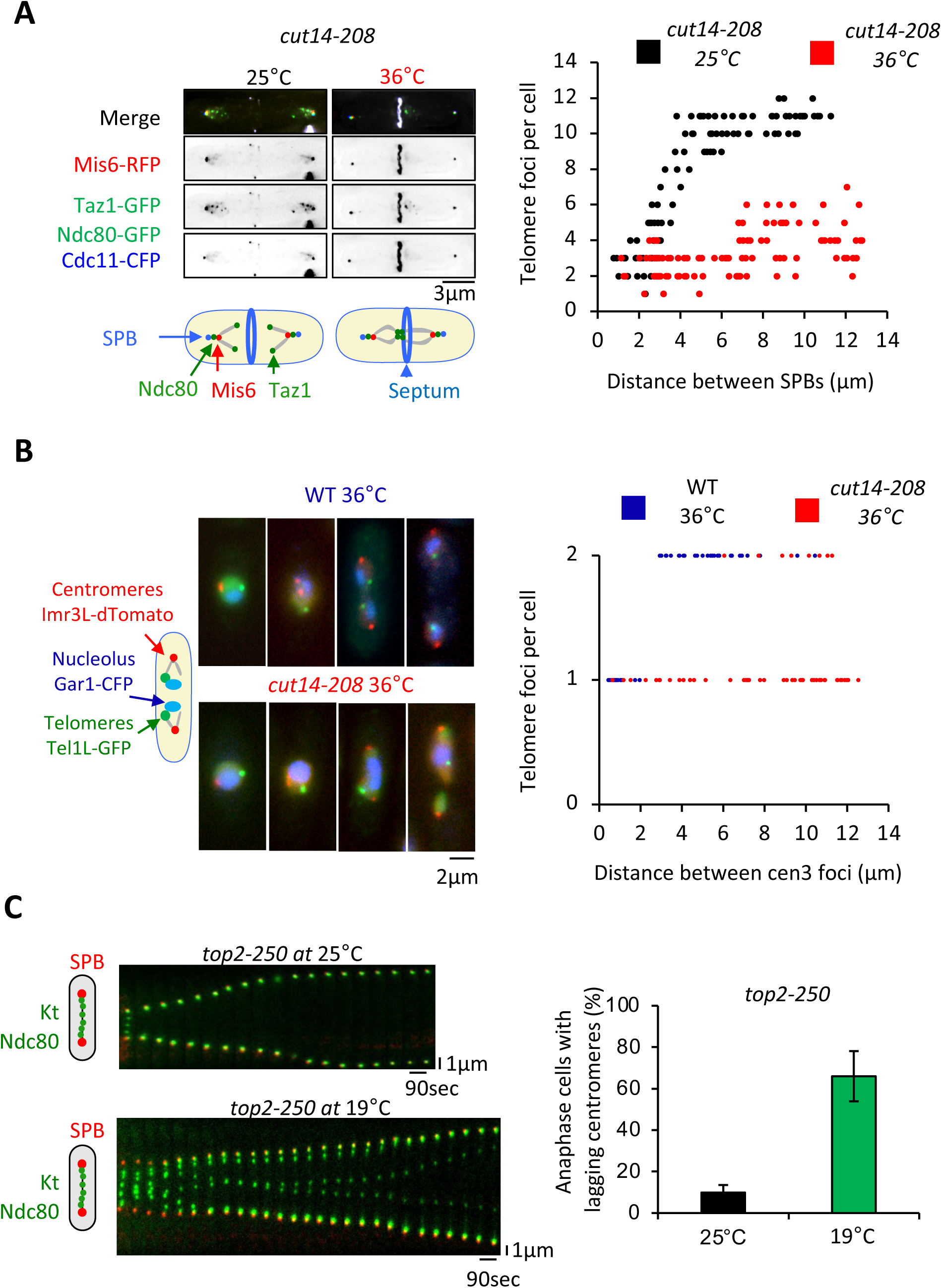
Condensin takes part in telomere disjunction during anaphase in a decatenation- independent manner. (**A**) left panel: Condensin mutant cells *cut14-208* grown at the permissive temperature (25°C) or shifted to the restrictive temperature of 36°C for three hours were fixed with formaldehyde and stained with calcofluor to reveal the septum. Telomeres were visualized via Taz1-GFP (green), kinetochores/centromeres via the colocalization of Mis6-RFP (red) and Ndc80- GFP (green), and spindle pole bodies (SPBs) via Cdc11-CFP (blue). Right panel: number of telomeric foci according to the distance between SPBs at 25°C and 36°C in the *cut14-208* mutant (n>90 cells for each strain). The data shown are from a single representative experiment out of three repeats. (**B**) Left panel: WT or *cut14-208* condensin mutant cells shifted to the restrictive temperature of 36°C for three hours were fixed with formaldehyde and directly imaged. Sister telomeres 1L (Tel1-GFP, green), sister centromeres 3L (imr3-tdTomato, red) and the nucleolus (Gar1-CFP) were visualized. Right panel: number of telomeric foci according to the distance between sister-centromeres 3L at 36°C (n>43 cells for each strain). The data shown are from a single representative experiment out of three repeats. (**C**) Kymograph representation of kinetochore dynamics in live *top2-250* mutant cells at permissive temperature (25°C) or restrictive temperature (18°C). Kinetochores are visualized via Ndc80-GFP (green) and SPBs via Cdc11- CFP (red). Right panel. Quantification of the percentage of *top2-250* anaphase cells showing lagging centromeres. Error bars indicate SD obtained from three independent experiments (n=100 for each condition).

**Figure S3.**
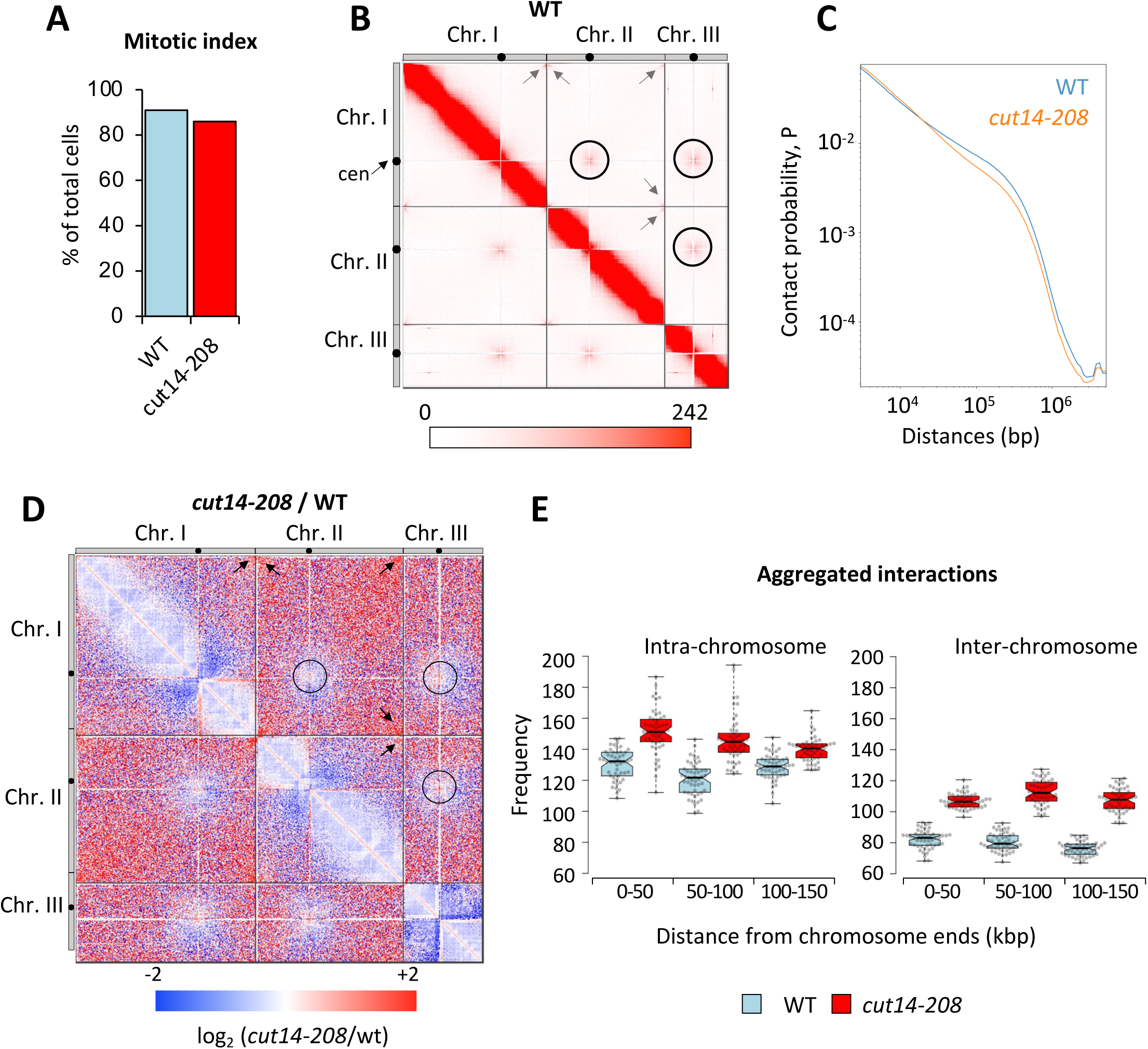
Condensin deficiency increases contact frequencies between telomeres in metaphase. Data from the biological and technical replicate of those shown in Fig. 3. (**A**) Mitotic indexes of cell cultures used for Hi-C. (**B**) Hi-C contact probability matrix at 25 kb resolution of wild-type cells arrested in metaphase at 33°C. Contacts between telomeres (arrows) and centromeres (circles) are indicated (**C**) Median contact probabilities as a function of distance along the chromosomes for wild-type and *cut14-208* cells arrested in metaphase at 33°C. (**D**) Hi-C difference map at 25 kb resolution comparing *cut14-208* and wild-type cells. (**E**) Aggregation of contact frequencies at chromosome ends in wild-type and *cut14-208* mutant cells in metaphase at 33°C. Boxes indicate the median, 1st and 3rd quartiles, whiskers extend to minimum and maximum values and notches represent the 95% confidence interval for each median. Data points are shown as grey circles.

**Figure S4.**
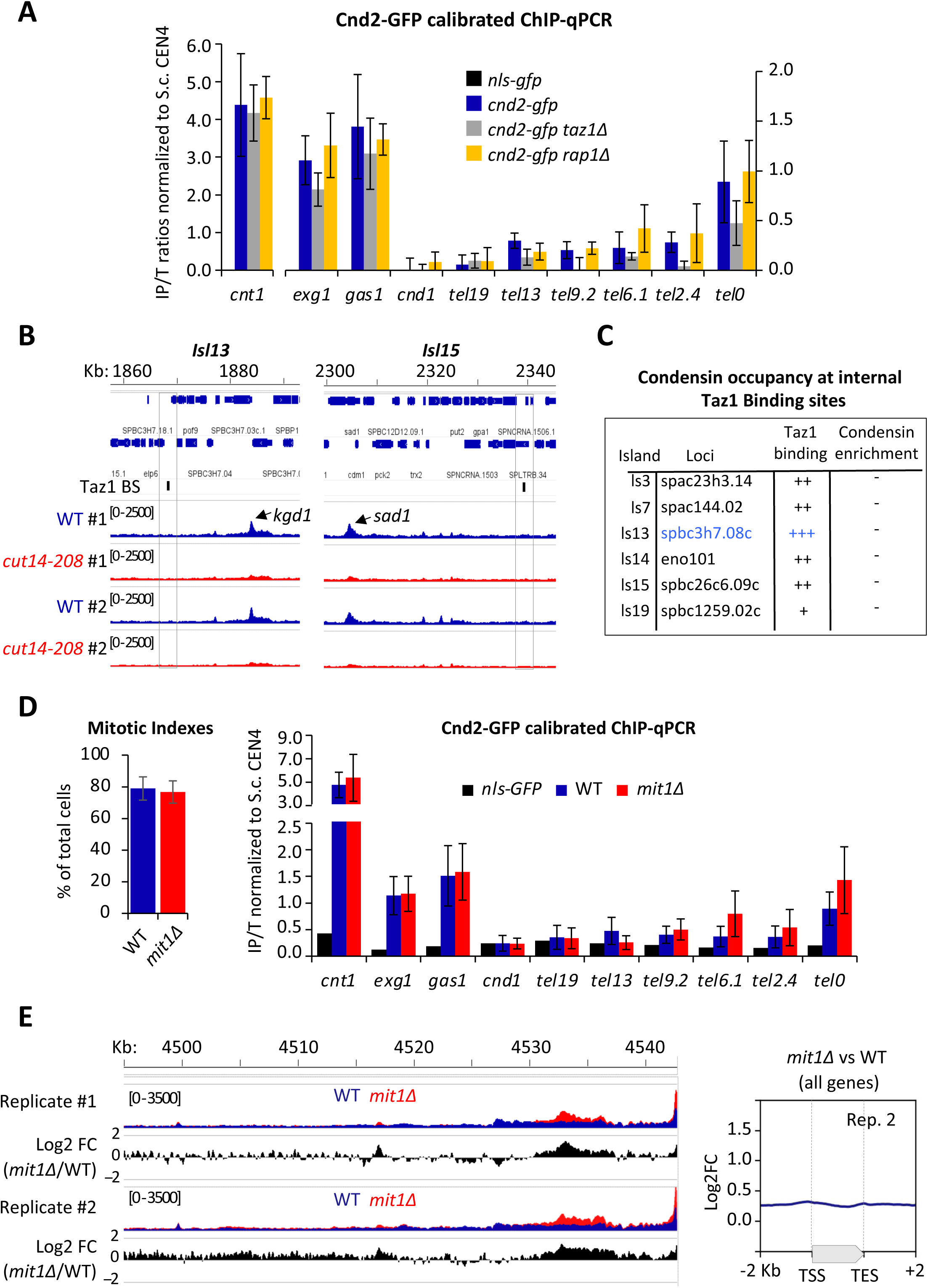
Taz1 specifically enriches condensin at telomeres. **(A)** Cnd2-GFP calibrated ChIP-qPCR shown in Figure 1C were normalized with respect to their corresponding IP/T ratios of budding yeast SMC3-GFP at CEN4. *cnt1* is the kinetochore domain of *cen1* while *exg1*, *gas1* and *cnd1* loci are high or low occupancy binding sites on chromosome arms. Shown are the average and standard deviations from n=3 biological and technical replicates. (**B**) Cnd2-GFP occupancy at non-telomeric Taz1 islands in wild-type or *cut14-208* mutant cells arrested in metaphase, as determined by calibrated ChIP-seq (n =2 biological and technical replicates). (**C**) Summary of the ChIP-seq results obtained at several representative Taz1 islands (Zofall *et al*, 2016). (**D**) Cnd2- GFP calibrated ChIP-qPCR from indicated metaphase arrests. Left panel: mitotic indexes of cell cultures used for ChIP. Right panel: results of calibrated ChIP-qPCR. Shown are averages and sd from 6 independent biological and technical replicates. (**E**) Left panel: genome browser views of Cnd2-GFP calibrated ChIP-seq in indicated metaphase arrests, with two independent biological and technical replicates (#). Replicate #1 is shown in Figure 4. Right panel: metagene profile of all condensin binding sites along chromosome arms from replicate #2. TSS (Transcription Start Site), TES (Transcription End Site).

**Figure S5.**
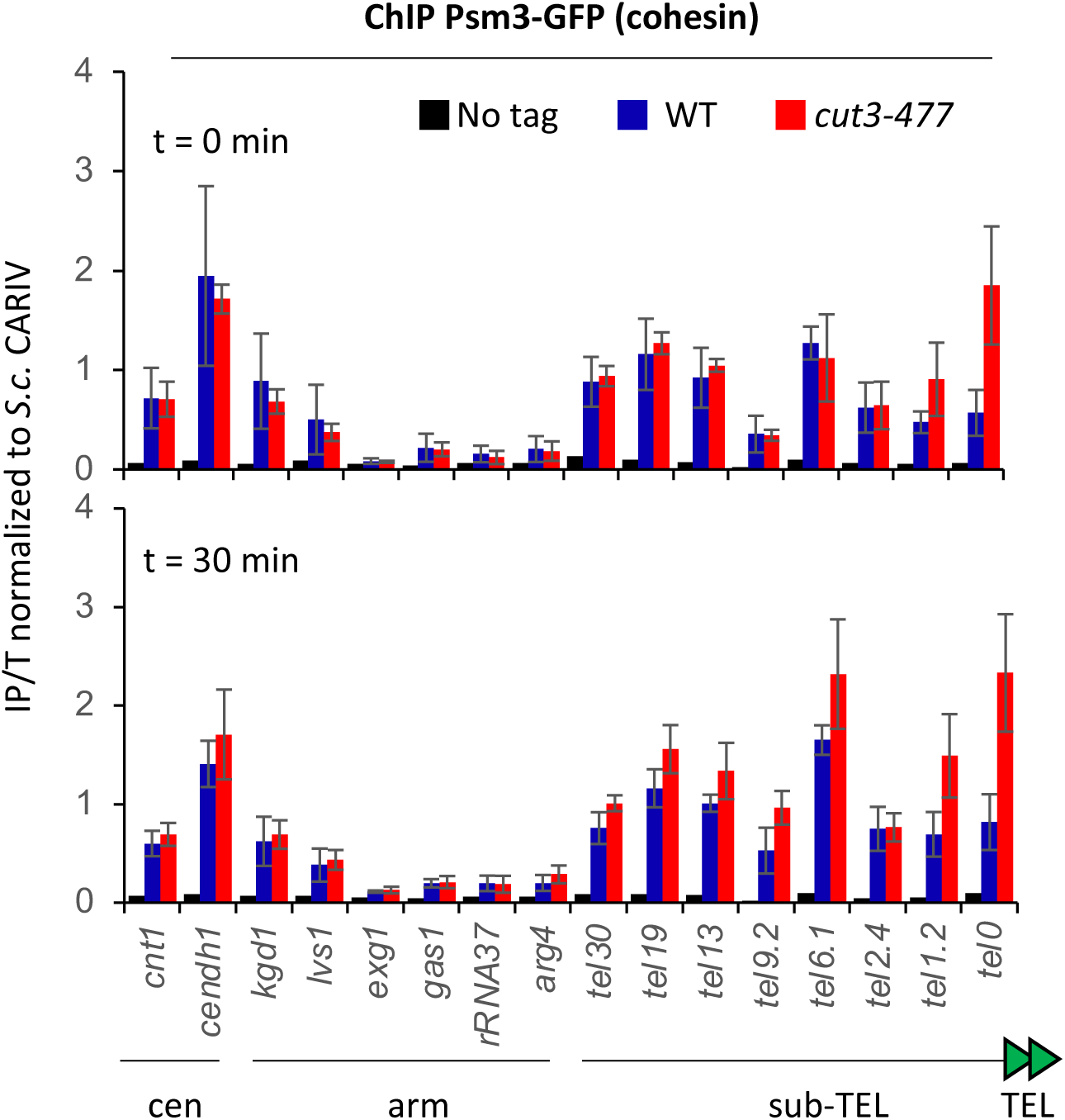
Condensin counteracts cohesin at telomeres. Psm3-GFP calibrated ChIP-qPCR from cells synchronized at G2/M (time 0 min; top panel) and upon their release in anaphase (time 30 min; bottom panel). *cendh1*, *kgd1* and *lvs1* loci are cohesin binding sites, while *exg1*, *gas1* and *rRNA37* loci are condensin binding sites. Percentage of IP with Psm3-GFP has been normalized using *S. cerevisiae* (S.c.) CARIV locus. Shown are the averages and standard deviations (sd) from 3 independent biological and technical replicates.

**Figure S6.**
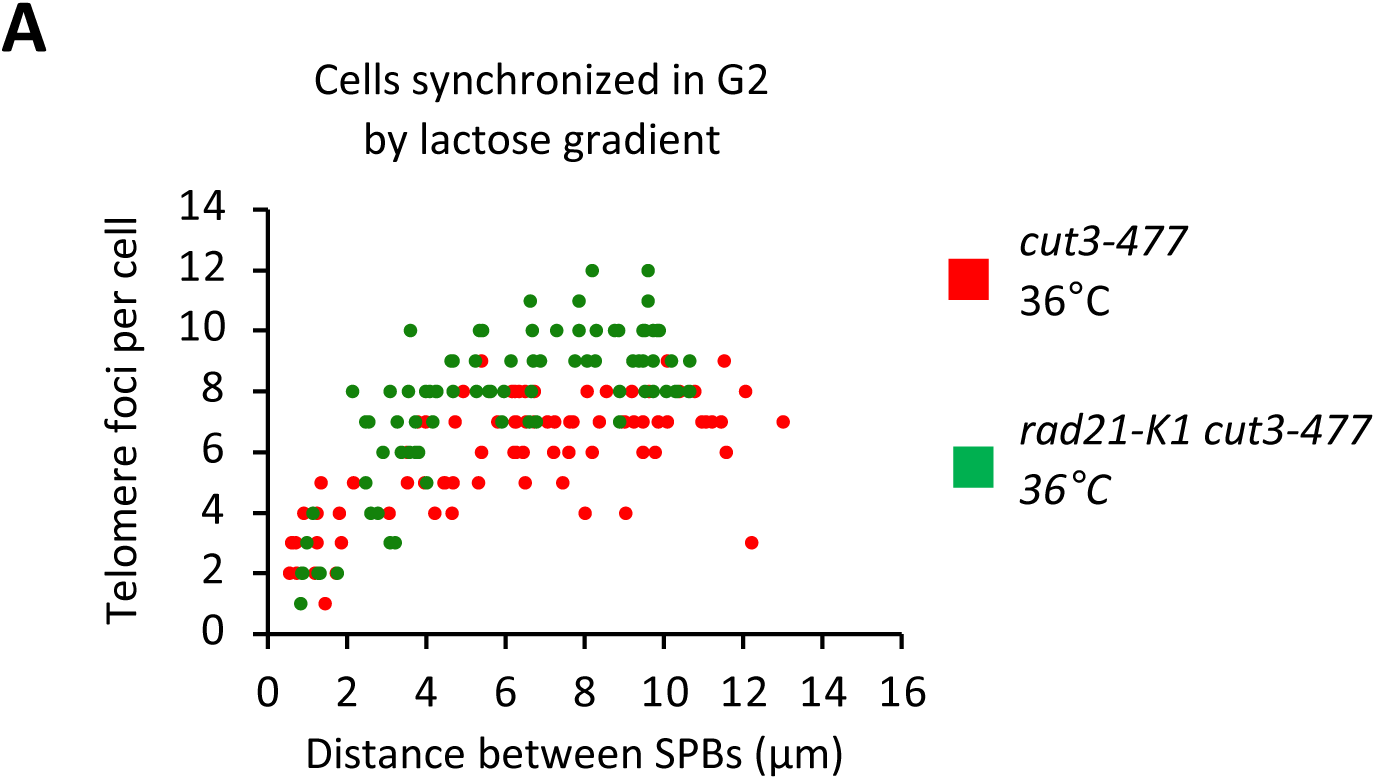
Condensin counteracts cohesin at telomeres. *cut3-477* and *cut3-477 rad21-K1* mutants were grown at permissive temperature and small, early G2 cells were purified using a lactose gradient. After synchronization, the entire cell population was in G2 (0% of cells in mitosis or cytokinesis). Purified early G2 cells were shifted to the restrictive temperature of 36°C and telomeric foci were scored according to the distance between SPBs (n>80 cells for each strain). The data shown are from a single representative experiment out of three repeats.

## DATA AND MATERIALS AVAILABILITY

Raw and processed data from calibrated-ChIP-seq, Hi-C and the TEL2R-extended ASM294v2 version of the fission yeast genome, are available at the NCBI Gene Expression Omnibus (GEO) repository under the accession number GSE196149.

## MATERIALS AND METHODS

### Media, molecular genetics and cell culture

Media, growth, maintenance of strains and genetic methods were as described (Moreno *et al*, 1991). Standard genetics and PCR-based gene targeting method (Bahler *et al*, 1998) were used to construct *S. pombe* strains. All fluorescently tagged proteins used in this study are expressed from single-copy genes under the control of their natural promoters at their native chromosomal locations. Strains used in this study are listed in Table S1. For metaphase arrests used in ChIP and Hi-C experiments, cells expressing the APC/C co-activator Slp1 under the thiamine-repressible promoter *nmt41* (Petrova *et al*, 2013) were cultured in synthetic PMG medium at 30°C, arrested in metaphase for 2h at 30°C by the adjunction of thiamine (20 µM final) and shifted at indicated restrictive temperatures for 1 hour. For Hi-C, the cultures were arrested for 3h at 33°C. Mitotic indexes were determined by scoring the percentage of cells exhibiting Cnd2-GFP fluorescence in their nucleoplasm (Sutani *et al*, 1999). G2/M block and release experiments were performed using an optimized cdc2-as allele (Aoi *et al*, 2014). Cells were arrested in late G2 by 3h incubation in the presence of 3-Br-PP1 at 2 µM final concentration (#A602985, Toronto Research Chemicals). Cells were released into synchronous mitosis by filtration and 3 washes with prewarmed liquid growing medium. For the viability spot assay, cell suspensions of equal densities were serially diluted five-fold and spotted on solid YES medium, the first drop containing 10^7^ cells. For microscopy, cells were grown in yeast extract and centrifuged 30 sec at 3000 g before mounting onto an imaging chamber. Total protein extractions for western blotting were performed by precipitation with TCA as previously described (Grallert & Hagan, 2017).

### Lactose gradient for G2 cells purification

Cell synchrony was achieved by lactose gradient size selection. Log phase cells (50ml of 5.10^6^ cells) were concentrated in 2ml and loaded onto a 50ml 7-35% linear lactose gradient at 4°C. After 10min centrifugation at 1600rpm at 4°C, 3ml of the upper of two visible layers was collected and washed twice in cold YES media before being resuspended in fresh medium. At this stage, cells were released at 36°C and fixed every 20min until the first mitotic peak.

### Cell imaging and fast microfluidic temperature control experiments

Live cell analysis was performed in an imaging chamber (CoverWell PCI-2.5, Grace Bio-Labs, Bend, OR) filled with 1 ml of 1% agarose in minimal medium and sealed with a 22 × 22-mm glass coverslip. Time-lapse images of Z stacks (maximum five stacks of 0.5 μm steps, to avoid photobleaching) were taken at 30 or 60 sec intervals. Images were acquired with a CCD Retiga R6 camera (QImaging) fitted to a DM6B upright microscope with a x100 1.44NA objective, using MetaMorph as a software. Intensity adjustments were made using the MetaMorph, Image J, and Adobe Photoshop packages (Adobe Systems France, Paris, France). Fast microfluidic temperature control experiments were performed with a CherryTemp from Cherry Biotech. To determine the percentage of chromatin bridges with unseparated telomeres, cells were fixed in 3.7% formaldehyde for 7 min at room temperature, washed twice in PBS, and observed in the presence of DAPI/calcofluor.

### Image processing and analysis

The position of the SPBs, kinetochores/centromeres and telomeres were determined by visualization of the Cdc11–CFP, Ndc80–GFP/Mis6-RFP and Taz1-GFP/Ccq1-GFP signals. Maximum intensity projections were prepared for each time point, with the images from each channel being combined into a single RGB image. These images were cropped around the cell of interest, and optional contrast enhancement was performed in MetaMorph, Image J or Photoshop where necessary. The cropped images were exported as 8-bit RGB-stacked TIFF files, with each frame corresponding to one image of the time-lapse series. For both channels, custom peak detection was performed. The successive positions of the SPBs were determined. The number of telomeres during spindle elongation was determined by visual inspection.

### Telomere length analysis by Southern blotting

Genomic DNA was prepared and digested with *Apa*I. The digested DNA was resolved in a 1.2% agarose gel and blotted onto a Hybond-XL membrane. After transfer, DNA was crosslinked to the membrane with UV and hybridized with a radiolabelled telomeric probe. The telomeric DNA probe was extracted by digestion of pIRT2-Telo plasmid by *Sac*I/*Pst*I.

### ChIP and calibrated-ChIP

Fission yeast cells, expressing either Cnd2-GFP or NLS-GFP, and arrested in metaphase by the depletion of Slp1, were fixed with 1% formaldehyde for 5 min at culture temperature and 20 min at 19°C in a water bath, quenched with glycine 0.125 M final, washed twice with PBS, frozen in liquid nitrogen and stored at -80°C until use. For calibration, *Saccharomyces cerevisiae* cells expressing Smc3-GFP were grown in Yeast Peptone Dextrose liquid medium at 30°C in log phase and fixed with 2.5% formaldehyde for 25 min. 2.10^8^ fission yeast cells were used per ChIP experiment. To perform calibrated ChIP the same amount of fission yeast cells was mixed with 4.10^7^ budding yeast cells. Cells were resuspended in lysis buffer (50 mM Hepes KOH pH 7.5, NaCl 140 mM, EDTA 1 mM, Triton X-100 1%, sodium deoxycholate 0.1%, PMSF 2mM) supplemented with a protease inhibitor cocktail (cat. 11836170001, Roche), and lysed with Precellys ®. Chromatin was sheared to ∼ 300 bp fragments with Covaris ® S220 (18 min at duty factor 5%, 200 cycles per burst, and 140W peak power), clarified twice by centrifugation at 9600 g at 4°C and adjusted to 1 mL final with lysis buffer. For ChIP, two 60 µl aliquots of chromatin each served as Total (input) fractions, while two aliquots of 300 µl of chromatin (IPs) were incubated each with 35 µl of Dynabeads^TM^ protein A (cat. 10002D, Invitrogen) and 8 µg of anti-GFP antibody (cat. A111-22, Invitrogen). For calibrated-ChIP-seq one 60 µl aliquot of chromatin served as Total (T) fraction and IP was performed on 600 µl of chromatin using 75 µl of Dynabeads^TM^ proteinA and 16 µg of anti-GFP antibody. T and IP samples were incubated overnight in a cold room, IPs being put on slow rotation. IPs were washed on a wheel at room temperature for 5 min with buffer WI (Tris-HCl pH8 20 mM, NaCl 150 mM, EDTA 2 mM, Triton X-100 1%, SDS 0.1%), WII (Tris-HCl pH8 20 mM, NaCl 500 mM, EDTA 2 mM, Triton X-100 1%, SDS 0.1%) and WIII (Tris-HCl pH8 10 mM, sodium deoxycholate 0.5%, EDTA 1 mM, Igepal 1%, LiCl 250 mM) and twice with TE pH8 without incubation. Immunoprecipitated materials on beads and T samples were brought to 100 µl in TE pH8, supplemented with RNAse A at 1µg/µl and incubated 30 min at 37°C. 20 µg of proteinase K was added and tubes were incubated 5h at 65°C. For calibrated-ChIPseq, IPs on beads were eluted in Tris 50 mM, EDTA 10 mM, SDS 1% 15 min at 65°C. Supernatants were transferred to a new tube supplemented with RNAse A at 1µg/µl and incubate 1h at 37°C. 200 µg of proteinase K was added followed by an incubation of 5h at 65°C. DNA was recovered using QIAquick PCR Purification Kit, following manufacturer’s instructions. For calibrated ChIP-qPCR, real time quantitative (q) PCRs were performed on a Rotor-Gene PCR cycler (Qiagen) using Quantifast (Qiagen) SYBR Green. The ratios (IP/T) calculated for fission yeast DNA sequences where normalized to their associated IP/T ratio calculated for budding yeast CARIV or CEN4 DNA sequences bound by SMC3-GFP. For calibrated ChIP-seq, Total and IPed DNA samples were washed with TE pH8 and concentrated using Amicon® 30K centrifugal filters, and libraries were prepared using NEBNext® Ultra™ II DNA Library Prep Kit for Illumina® kits according to the manufacturer’s instructions. DNA libraries were size-selected using Ampure XP Agencourt beads (A63881) and sequenced paired-end 150 bp with Novaseq S6000 (Novogene®).

### Hi-C sample preparation

Fission yeast cells, expressing Cnd2-GFP and arrested in metaphase by the depletion of Slp1 were fixed with 3% formaldehyde for 5 min at 33°C followed by 20 min at 19°C, washed twice with PBS, frozen in liquid nitrogen and stored at -80°C. 2.10^8^ cells were lysed in ChIP lysis buffer with Precellys ®. Lysates were centrifuged 5000 g at 4°C for 5 min and pellets were resuspended once in 1 ml lysis buffer and twice in NEB^®^ 3.1 buffer. SDS was added to reach 0.1% final and samples were incubated for 10 min at 65°C. SDS was quenched on ice with 1% Triton X-100 and DNA digested overnight at 37°C with 200 Units of *Dpn*II restriction enzyme. Samples were incubated at 65°C for 20 min to inactivate *Dpn*II. Restricted-DNA fragments were filled-in with 15 nmol each of biotin-14-dATP (cat. 19524016, Thermofisher), dTTP, dCTP and dGTP, and 50 units of DNA Klenow I (cat. M0210M, NEB) for 45 min at 37°C. Samples were diluted in 8 ml of T4 DNA ligase buffer 1X and incubated 8 hours at 16°C with 8000 Units of T4 DNA ligase (NEB). Crosslinks were reversed overnight at 60°C in the presence of proteinase K (0.125 mg / ml final) and SDS 1% final. 1 mg of proteinase K was added again and tubes were further incubated for 2 hours at 60°C. DNA was recovered by phenol-chloroform-isoamyl-alcohol extraction, resuspended in 100 µl TLE (Tris/HCl 10 mM, 0.1 mM EDTA, pH8) and treated with RNAse A (0.1 mg / ml) for 30 min at 37°C. Biotin was removed from unligated ends with 3 nmol dATP, dGTP and 36 Units of T4 DNA polymerase (NEB) for 4 hours at 20°C. Samples were incubated at 75°C for 20 min, washed using Amicon® 30k centrifugal filters and sonicated in 130 µl H2O using Covaris® S220 (4 min 20°C, duty factor 10%, 175W peak power, 200 burst per cycle). DNA was end-repaired with 37.5 nmol dNTP, 16.2 Units of T4 DNA polymerase, 54 Units of T4 polynucleotide kinase, 5.5 Units of DNA Pol I Klenow fragment for 30 min at 20°C and then incubated for 20 min at 75°C. Ligated junctions were pulled-down with Dynabeads® MyOne™ Streptavidin C1 beads for 15 min at RT and DNA ends were A-tailed with 15 Units of Klenow exo- (cat. M0212L, NEB). Barcoded PerkinElmer adapters (cat. NOVA-514102) were ligated on fragments for 2 hours at 22°C. Libraries were amplified with NextFlex PCR mix (cat. NOVA- 5140-08) for 5 cycles, and cleaned up with Ampure XP Agencourt beads (A63881). Hi-C libraries were paired-end sequenced 150bp on Novaseq6000.

### Calibrated ChIP-seq data analysis

Scripts and pipelines are available in the git repository https://gitbio.ens-lyon.fr/LBMC/Bernard/chipseq (tag v0.1.0). Analyses have been performed based on the method described by Hu et al. (Hu *et al*, 2015), using a modified version of the nf-core/chipseq (version 2.0.0) pipeline (https://doi.org/10.1038/s41587-020-0439-x) executed with nextflow (version 23.02.1). We used the *S. pombe* genome (ASM294v2, or its TEL2-R extended version available at Omnibus GEO GSE196149) and the *S. cerevisiae* genome (sacCER3 release R64-1-1).) for internal calibration. Technical details regarding our calibrated-ChIP-seq pipeline are available in the Appendix Supplementary Methods section. Of note, we noticed a sharp decrease in the number of reads passed the coordinate 4,542,700 at the right end of chromosome II, i.e. within the telomeric repeats of TEL2R, in Total extracts. Thus, to avoid any biased enrichment in our IP/Total ratios, all calibrated ChIP-seq results concerning TEL2R have been taken within the limit of the position 4,542,700 within telomeric repeats of TEL2R.

### Hi-C data analysis

Computational analyses of Hi-C data were performed with R (version 3.4.3). Reads were aligned on the genome of *S. pombe* version ASM294v2 using bwa (version 0.7.17-r1188) with default settings. Hi-C contacts matrices of DpnII digested genomic fragments were normalized and processed using Juicer (version 1.6: https://github.com/aidenlab/juicer). Hi-C reads were binned to a resolution of 5 kb using a square root vanilla count normalization. 2D plots were performed for normalized (observed/expected) Hi-C read counts using Juicebox. Differential 2D plots were visualized in Log2 (of normalized Hi-C reads in mutants / normalized Hi-C reads in wild-type control). Aggregation of Hi-C data was performed essentially as previously described (Liang *et al*, 2014; Rao *et al*, 2014) with the following modified parameters for adaptation to 3D contacts at telomeres: Hi-C reads were counted over bins of 5 kb over a 150 kb distal region covering both telomeres of each chromosome. Long-range interactions were assessed for all combinations of telomeres over a sliding matrix (21x21 bins). To optimize detection of long-range interactions between telomeres, a quantilization was performed by ranking the 21x21 bins of every sub-matrix contributing to the aggregation, allowing to assess interactions from averaged values. Statistical analyses were performed both for the corresponding quantilized matrices, and verified with non- quantilized matrices, using a Mann-Whitney-Wilcoxon test using R (Stats4) and validated for each of the replicates made for every mutant and wild-type conditions.

**Table S1.** Strain list used in this study.

| Strain | Genotype | Figures |
| --- | --- | --- |
| LY4682 | *h- leu1-32 ura4D18 cdc2asM17* | Fig. 1B |
| LY4731 | *h- leu1-32 ura4D cdc2asM17 cnd2-GFP-LEU2* | Fig. 1B |
| LY6281 | *Mata ade2-1 his3-11 his3-15 ura3 leu2-3 trp1-1 can1-100 SMC3-GFP-KanR* | Fig. 1C-D, S1, 4 & S4 |
| LY4483 | *h- leu1-32 ura4D-18 ura4-Pnmt41-slp1+ cnd2-GFP-LEU2* | Fig. 1C-E, S1, 3 & S3, 4 & S4 |
| LY6305 | *h- leu1-32 ura4D cut14-208 ura4-Pnmt41-slp1+ cnd2-GFP-LEU2* | Fig. 1C-E, S1D, 3 & S3, S4B |
| LY6304 | *h+ leu1-32 ura4D cut3-477 ura4-Pnmt41-slp1+ cnd2-GFP-LEU2* | Fig. 1E, S1E |
| LY5480 | *leu1- ura4- ade6- ura4-Pnmt41-slp1+ aur1::padh21-NLS-GFP-9PK::aur1R* | Fig. 4A, 4C, S4A & S4D |
| LY5298 | *h? leu1-32 ura4D18 ade6-21? KanR-Pnmt1-slp1+ taz1Δ::ura4+ cnd2-GFP-LEU2* | Fig. 4A, 4C, S4A |
| LY5948 | *h- leu1-32 ura4D ura4-Pnmt41-slp1+ rap1Δ::KanR cnd2-GFP-LEU2* | Fig. 4A, 4C, S4A |
| LY5898 | *h+ leu1-32 ura4D ura4-Pnmt41-slp1+ mit1Δ::KanR cnd2-GFP-LEU2* | Fig. 4D-E, S4D-E |
| ST1711 | *h+ cdc11-CFP::KanR taz1-GFP::KanR mis6-2mRFP::hph* | Fig. 2A-B |
| ST2624 | *h?cut3-477 cdc11-CFP::KanR taz1-GFP::KanR mis6-2mRFP::hph* | Fig. 2B |
| ST1738 | *h? cut14-208 cdc11-CFP::KanR taz1-GFP::KanR mis6-2mRFP::hph* | Fig2A & Fig.S2A |
| ST2622 | *h?cut3-477 cdc11-CFP::KanR taz1-GFP::KanR* | Fig.2D, 5C, 6A |
| ST1665 | *h? top2-250 cdc11-CFP:: KanR taz1-GFP:: KanR* | Fig.2D-E |
| FY24566 | *h+ imr3L-tetO-ura4+ Z:adh31-tetR-tdTomato<<natr his7+<<Pdis1-GFP-lacI leu1 ade6 tel1L-lacO-kanr C:adh21-gar1-CFP<<hygr (Tada et al. 2011)* | Fig.S2B |
| ST2640 | *h? cut14-208 imr3L-tetO-ura4+ Z:adh31-tetR-tdTomato<<natr his7+<<Pdis1-GFP-lacI leu1 ade6 tel1L-lacO-kanr C:adh21-gar1-CFP<<hygr* | Fig.S2B |
| PR109 | *h- leu1-32 ura4D-18* | Fig. 5B, S2C |
| SC388 | *h- leu1-32 ura4D-18 ade6-M210 taz1 ::ura4+* | Fig. 5B |
| LY999 | *h- leu1-32 ura4D ade6-M210 cut3-477 :NatR* | Fig. 5B, S2C |
| SC1432 | *h ? leu1-32 ura4D-18 taz1::ura4+ cut3-477 :NatR* | Fig. 5B |
| ST1928 | *h? taz1::ura+ cut3-477 ccq1-GFP::KanR cdc11-CFP:: KanR ura4D-18* | Fig. 5A |
| ST1664 | *h? top2-250 ndc80-GFP:: KanR cdc11-CFP:: KanR mis6-RFP-hyg* | Fig. S2C |
| ST1172 | *h? ccq1-GFP:: KanR cdc11-CFP:: KanR* | Fig. 5A |
| ST1183 | *h- taz1::KanR ccq1-GFP::KanR cdc11-CFP:: KanR* | Fig. 5A |
| ST1657 | *h- cut3-477 ccq1-GFP:: KanR cdc11-CFP :: KanR* | Fig. 5A |
| ST2636 | *h? mit1::hph cdc11-CFP:: KanR taz1-GFP:: KanR* | Fig. 5C |
| LY646 | *h- leu1-32 ura4D-18 cut14-90* | Fig. 2C |
| LY140 | *h- leu1-32 ura4D cut14-208* | Fig. 2C |
| LY1252 | *h- leu1-32 ura4D ade6-M210 cut14-180* | Fig. 2C |
| LY3185 | *h- ura4D ade6-M210 cut3-M26* | Fig. 2C |
| LY3186 | *h- ura4D ade6-M210 cut3-I23-GFP:his7+* | Fig. 2C |
| LY4681 | *h+ leu1-32 ura4D18 cdc2asM17* | Fig. 6B, S5 |
| LY7260 | *h+ leu1-32 ura4D? ade6-210 cdc2asM17 psm3-GFP-NatR* | Fig. 6B, S5 |
| LY7262 | *h+ leu1-32 ura4D? ade6-210 cdc2asM17 psm3-GFP-NatR cut3-477* | Fig. 6B, S5 |
| STCS138 | *h? rad21-K1 cdc11-CFP:: KanR taz1-GFP:: KanR* | Fig.6A, S6A |
| ST2638 | *h? rad21-K1 cut3-477 cdc11-CFP:: KanR taz1-GFP:: KanR* | Fig.6A, S6A |

**Table S2.**
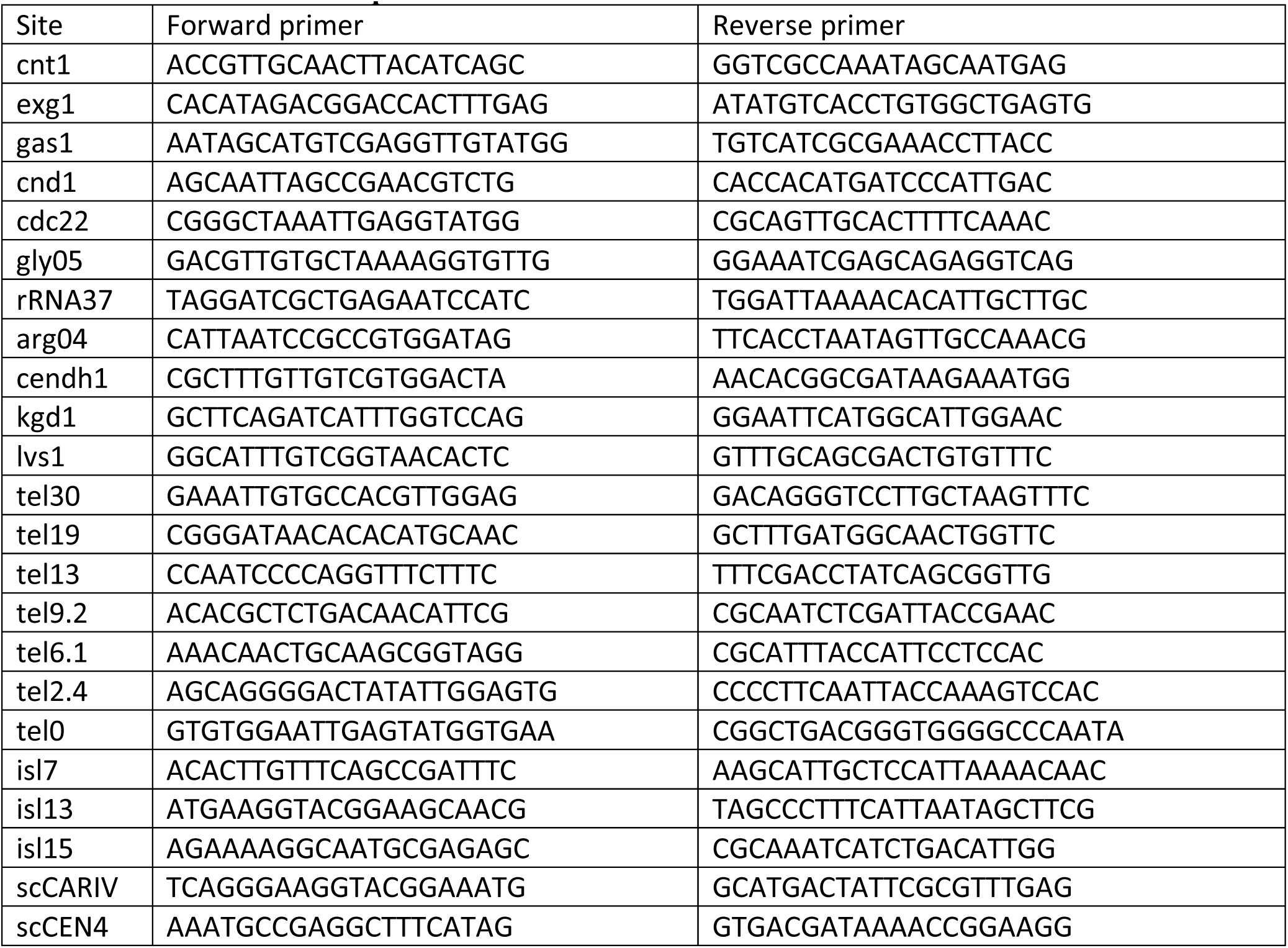
Primers used for qPCR.

**Table S3.** Antibodies used in this study.

| Antibody | target | experiment |
| --- | --- | --- |
| A-11122 (Invitrogen) | Anti-GFP | ChIP |
| Tat-1 (a gift from Keith Gull) | Anti-α-Tubulin | Western blot |
| Cnd2 (euromedex, GTX64102) | Anti-Cnd2 fission yeast | Western blot |
| anti-rabbit NA9340V Amersham | Anti-rabbit secondary | Western blot |
| anti-mouseNA931 -1ml Amersham | Anti-mouse secondary | Western blot |

## Appendix Supplementary Methods

Scripts and pipelines are available in the git repository https://gitbio.ens-lyon.fr/LBMC/Bernard/chipseq (tag v0.1.0). Pipelines were executed with nextflow (version 23.02.1). In the subsequent section, *genome* refers to the *S. pombe* genome (*ASM294v2, or its TEL2-R extended version available at* Omnibus GEO GSE196149), while *calibration genome* refers to the genome of *S. cerevisiae* (sacCER3 release R64-1-1) used for internal calibration. To perform the analyses, we use a modified version of the nf-core/chipseq (version 2.0.0) pipeline (https://doi.org/10.1038/s41587-020-0439-x). We modified this pipeline as follows: We added an optional `--fasta_calibration` parameter to pass the calibration genome. We modified the subworkflow `prepare_genome.nf` to run `GUNZIP_FASTA` on both the `--fasta` and `-- fasta_calibration` files and merge the two fasta file with the `MIX_FASTA` process, while adding a `cali_` prefix to the names of the chromosome of the calibration genome. The workflow chipseq.nf was modified to run a new process, `BAM_CALIB` (replacing `BEDTOOLS_GENOMECOV`), on the IP bam file and their corresponding Total (INPUT) bam file to generate calibrated IP and INPUT bigwig files. The workflow chipseq.nf was modified to then run `BIGWIG2BAM` to generate synthetic bam files from the output of `BAM_CALIB.` These synthetic bam files are single-end and replace the output of the mapping step in the next parts of the pipeline. The `BIGWIG2BAM` outputs are indexed with a `SAMTOOLS_INDEX` process. The `BAM_CALIB` tool (https://gitbio.ens-lyon.fr/LBMC/Bernard/bamcalib v0.1.5) takes into inputs a sorted bam file for the IP data and a sorted bam file for the TOTAL data (mapped on the concatenation of the two genomes), and output a calibrated bigwig for the reference genome. For the normalization, we modified a previously described method (Hu *et al*, 2015) in order to account for biases in coverages in the TOTAL fractions. We introduce the following notation: *IP*_*x*_(*t*) is the coverage at position *t* in the IP sample on the reference genome, *IP*_*c*_(*t*) is the coverage at position *t* in the IP sample on the calibration genome, *WCE*_*x*_(*t*) is the coverage at position *t* in the TOTAL (whole cell extract) sample on the reference genome, and *WCE*_*c*_(*t*) the coverage at position *t* in the TOTAL sample on the calibration genome. In a reference genome of size *T*_*x*_ (ignoring the chromosome segmentation), and in a calibration genome of size *T*_*c*_, Hu et al. compute the Occupancy Ratio (OR) as follows (Hu *et al*, 2015):

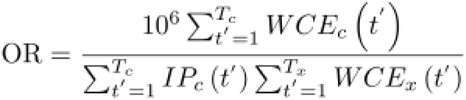

Instead, we used the following formula with *β* (default to 10^3^) an arbitrary scaling factor.

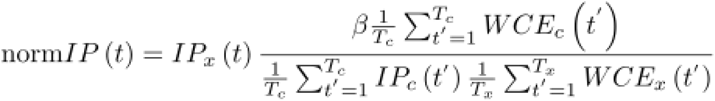

This formula can be described as follows:

The technical variations on the IP efficiencies are corrected by scaling IP*_x_*(*t*) by the calibration genome coverage:

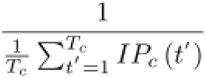

To account for variations in cells proportion, we correct by a scaled WCE coverage:

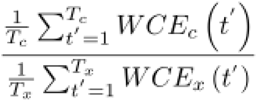

To be able to analyze the coverage information at repetitive regions of the genome, we propose to normalize the signal nucleotide by nucleotide and introduce the OR ratio:

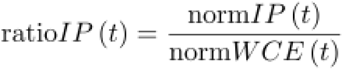

with:

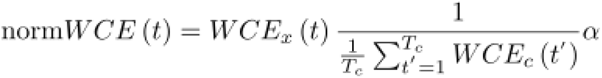

We then find *α* such that (not to distort the *IP* signal on average):

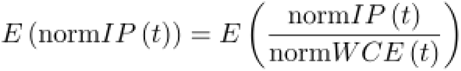

which gives

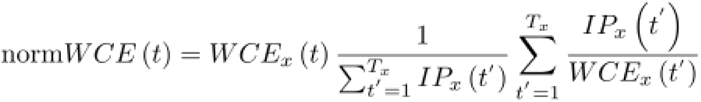

With this method, we retain the internal calibration developed by Hu et al. (Hu *et al*, 2015) and we account for variations in read density at each base in WCE samples.

In the BAM_CALIB tools, the coverage does not correspond to the number of read covering a given position like in classical tools outputting bigwig. Instead we compute the number of fragments for paired-end data. To compute the coverage density X_y_(*t*) with X ∈ [*IP*,*WCE*] and y ∈ [*c*,*x*] we count the number of reads *r*(*t*) overlapping with position . For properly paired reads (with a mate read on the same chromosome and with a starting position ending after the end of the read) we also count a density of 1 between the end of the first reads and the start of his mate read X_y_(*t*) = *r*(*t*)+*g*(*t*). Some fragment can be artificially long (with reads mapping to repeated regions at the start and end of a chromosome), therefore, we compute a robust mean of the gap size, between two reads of a pair, by removing the 0.1 upper and lower quantile of the fragment length distribution. Fragments with a size higher than *ϕ*^-1^ (0.95,*μ*,1,0) are set to end at the *ϕ*^-1^ (0.95,*μ*,1,0) value, with *ϕ*() the Normal CDF function. For fragments shorter than the read length, we don’t count the overlapping reads region as a coverage of 2 fragments but as the coverage of 1 fragment. The BIGWIG2BAM Tools (https://gitbio.ens-lyon.fr/LBMC/Bernard/bamcalib v0.1.1), generate a synthetic bam file from a bigwig file and a reference genome. The purpose of this tool is to create a bam file having the same coverage profile as the one described in the input bigwig file. Therefore, we can run any chip-seq tools working with bam file instead of bigwig file on our normalized data.

